# Evaluation of single-sample network inference methods for precision oncology

**DOI:** 10.1101/2023.07.11.548508

**Authors:** Joke Deschildre, Boris Vandemoortele, Jens Uwe Loers, Katleen De Preter, Vanessa Vermeirssen

## Abstract

A major challenge in precision oncology is to identify targetable cancer vulnerabilities in individual patients. Modelling high-throughput omics data in biological networks allows identifying key molecules and processes of tumorigenesis. Traditionally, network inference methods rely on many samples to contain sufficient information for learning and predicting gene interactions for a group of patients. However, to implement patient-tailored approaches in precision oncology, we need to interpret omics data at the level of the individual patient. Several single-sample network inference methods have been developed that infer biological networks for an individual sample from bulk RNA-seq data. However, only a limited comparison of these methods has been made. Moreover, many methods rely on ‘normal tissue’ samples as reference point for the tumor samples, which is not always available.

Here, we conducted an evaluation of the single-sample network inference methods SSN, LIONESS, iENA, CSN and SSPGI using expression profiles of lung and brain cancer cell lines from the CCLE database. The methods constructed networks with distinct network topologies, as observed by edge weight distributions and other network characteristics. Further, hub gene analyses revealed different degrees of subtype-specificity across methods. Single-sample networks were able to distinguish between tumor subtypes, as exemplified by edge weight clustering, enrichment of known subtype-specific driver genes among hub gene sets, and differential node importance. Finally, we show that single-sample networks correlate better to other omics data from the same cell line as compared to aggregate networks. Our results point to the important role of single-sample network inference in precision medicine.

## Introduction

In order to understand the complex molecular interactions at play in tumor pathogenesis, high-throughput omics data have been generated at an increasing pace^1^. Modeling these data in biological networks allows for determining the key molecules and processes that drive tumorigenesis^2, 3^. Traditionally, network inference methods rely on many samples to contribute sufficient information to the learning process and to counteract the curse of dimensionality in the omics data, i.e. the number of genes by far outnumbering the number of samples. Methods to accomplish this on tissue level from bulk omics data are already well-established, and rely on varying underlying statistical and mathematical principles such as correlation, mutual information, Bayesian networks and regression^4–6^. We and others have shown that different computational methods reveal complementary aspects of the ‘true’ underlying networks^4, 7–9^. However, these methods infer networks based on numerous samples and therefore determine a general estimate of gene interactions largely shared by that group of samples. Hence, they result in population-level networks, averaging the phenotypic effects of individual patients or samples. For clinical applications, we need to be able to interpret and extract meaningful information from omics data of a single individual to be able to direct individualized treatment in precision medicine^10^.

Currently, several approaches are being explored to analyze omics data from a single sample or patient, and are referred to in literature as single-sample, single-subject, sample-specific, patient-specific and personalized methodologies. Deep n-of-1 phenotyping, where multiple omics are profiled in a single individual at different locations in the body longitudinally, is envisioned to be essential for the early detection and personalized treatment of cancers^11^. Obtaining multiple samples from one patient is nonetheless not straightforward due to an increased cost, increased surgical risk, or limited tumor size. Moreover, single-sample or patient-specific networks can be built from single cell RNA-seq data of a single subject, where the profiling of many cells inherently contains the variability required to infer the statistical dependencies between genes^12^. However, single cell omics data have specific limitations such as high-dimensionality, sparsity and overdispersion and network inference methods are still being optimized to deal with these issues. Also, single cell technologies are currently more expensive and hardly implemented in the clinic as compared to bulk protocols. On bulk transcriptome data, several methods extract relevant biological knowledge from individual samples without requiring a large disease cohort, as reviewed in^13^. They either provide a gene-centric view on differentially expressed (DE) genes or a pathway-centric view on deregulated pathways, comparing a single sample against a reference cohort or a control sample^13–16^. In addition, VIPER can predict protein activity from regulon enrichment on single-sample gene expression signatures obtained using a reference set^17^. The single-sample Network Perturbation Assessment (ssNPA) is a method for subtyping samples based on single-sample deregulation of their gene networks^18^. While these methods allow for biological interpretation of omics data at the individual level, they do not generate biological networks or gene interactions for single samples or patients.

To address this, several single-sample network inference methods have been developed that can infer a biological network for a single sample of bulk RNA-seq data. Several of these methods make use of an aggregate network constructed from all samples and a statistical wrapper to infer single-sample features within these networks. Optionally, the aggregate networks can be pruned by a background network, e.g. an experimentally validated protein-protein interaction network. The Single-Sample Network (SSN) algorithm calculates the significant differential network between the Pearson Correlation Coefficient (PCC) networks of a set of reference samples on the one hand and that same reference set plus the sample of interest on the other hand, both using the STRING database as background network^19^. The authors experimentally validated that SSN identified functional driver genes contributing to resistance in non-small cell lung cancer cell lines. Subsequently, SSN has been applied to breast and colon cancer to study stage-and subtype-related networks and to identify diagnostic and prognostic biomarkers^20, 21^. LIONESS also uses a leave-one-out approach in aggregate network inference to come to a single-sample network, and through linear interpolation incorporates information on both the similarities and the differences between the networks with and without the sample of interest^22^. LIONESS has the major advantage that any network inference method of choice can be used to construct the aggregate networks, and has been applied e.g. to study sex-linked differences in colon cancer drug metabolism^23^. The Individual-specific Edge-Network Analysis (iENA) algorithm constructs single-sample PCC node-networks and single-sample higher-order PCC edge-networks by altered PCC calculations of the expression data of the sample of interest and a set of reference samples^24^. On the other hand, Sample Specific Perturbation of Gene Interactions (SSPGI) computes individual edge-perturbations based on differences between the rank of genes within the expression matrix of normal samples and individual samples of interest^25^. The Cell-Specific Network construction (CSN) method transforms the expression data into more stable, statistical gene associations, rendering a binary network output at single cell or single-sample resolution, for single or bulk RNA-seq data respectively^26^.

The above mentioned single-sample network inference methods have mainly been applied by the research groups that developed them and a systemic neutral comparison is still missing. Only limited comparisons have been performed, which either focused on a limited number of methods, focused on downstream network control methods or made use of metabolomics data with a limited number of features^22, 27–29^. Furthermore, many of the compared methods rely on ‘normal tissue’ reference samples to contrast the tumor samples to, which might not be available for all tumor types or in all precision oncology cases. The CCLE database offers multiple omics, including transcriptomics, on a large panel of comprehensively characterized human cancer models/cell lines and thus represents an ideal playground to apply and compare single-sample network inference methods^30, 31^. In this study we constructed single-sample networks using SSN, LIONESS, iENA, CSN and SSPGI for lung cancer and brain cancer samples. We found that each method constructed networks with distinct topologies at the level of edge weight distributions and network characteristics. Classification of tumor subtypes by edge weight clustering and hub gene analysis, which revealed different degrees of subtype-specificity across methods, showed us that SSN, LIONESS and iENA are most capable of emphasizing subtype-specific features. Differentially important nodes however were not enriched for subtype specific known driver genes. Finally, we show that single-sample networks correlated better to other omics data from the same cell line as compared to aggregate networks.

## Results

### Subtype-specific gene expression in lung and brain CCLE cell lines

In order to evaluate single-sample network inference methods in the absence of healthy control reference samples, we set out to compare SSN, LIONESS, iENA, CSN and SSPGI on gene expression profiles from CCLE lung and brain cancer cell lines^30^. We identified cell lines that closely matched their corresponding tumor tissue with regard to gene expression, and retained 86 lung and 67 brain cancer cell lines (Methods)^32^. These are further split into subtypes including 73 non-small cell lung carcinoma (NSCLC), 12 small cell lung carcinoma (SCLC), 1 lung carcinoid, 36 glioblastoma, 9 astrocytoma, 8 glioma, 9 medulloblastoma, 3 meningioma, 3 oligodendroglioma and 2 primitive neuroectodermal tumor (PNET) cell lines (Methods). An initial clustering of lung expression profiles showed that all but one of SCLC samples clustered separately from NSCLC samples (**Figure 1A**). We further compared gene expression in both cancer subtypes and identified 1510 up-and 1553 downregulated genes in NSCLC versus SCLC samples (absolute log fold change (abs(LFC)) >= 1, adjusted p-value (padj) <= 0.05) (**Figure 1B**). A clustering of brain expression profiles revealed one subcluster containing all but one of medulloblastoma and all PNET samples (**Figure 1C**). Due to limited sample sizes for meningioma, oligodendroglioma, PNET and glioma, we choose to perform subsequent differential analyses in brain between glioblastoma and medulloblastoma samples only. In total, 1354 and 1043 genes were up- and downregulated in glioblastoma versus medulloblastoma samples (**Figure 1D**). Hence, we detected substantial transcriptional differences between tumor subtypes for both lung and brain samples.

### Construction of single-sample networks

For both tumor types, we selected highly-variable genes (HVG) for network construction. First, we inferred an aggregate, undirected co-expression network using PCC, representing all samples. Next, single-sample networks were then inferred using LIONESS, SSN, iENA, SSPGI and CSN. We slightly modified several tools to run with PCC as the underlying network inference method and in absence of ‘normal tissue’ reference samples (Methods, GitHub). The choice of PCC as underlying network inference approach allowed for a consistent comparison between single-sample networks, as some methods exclusively function with PCC, and between the single-sample and the aggregate networks. We further pruned the aggregate and single-sample networks by selecting edges present in the HumanNet network, an integrated human functional gene network that was used as background network (Methods, **Figure 2**)^33^. A total of 5454 nodes and 53 296 edges were present in the lung aggregate HumanNet network and in single-sample lung HumanNet networks, covering respectively 30.50% of proteins and 10.14% of HumanNet interactions. Lung SSPGI networks were slightly smaller and comprised 4814 nodes connected by 43.193 edges (Methods, **Suppl. Table 1**). The aggregate brain network and single-sample brain networks constructed using SSN, LIONESS, CSN and iENA comprised 4741 nodes and 42 948 edges after pruning for HumanNet interactions, while 4686 nodes and 42.206 edges remained in the SSPGI networks after pruning.

**Figure 1.**
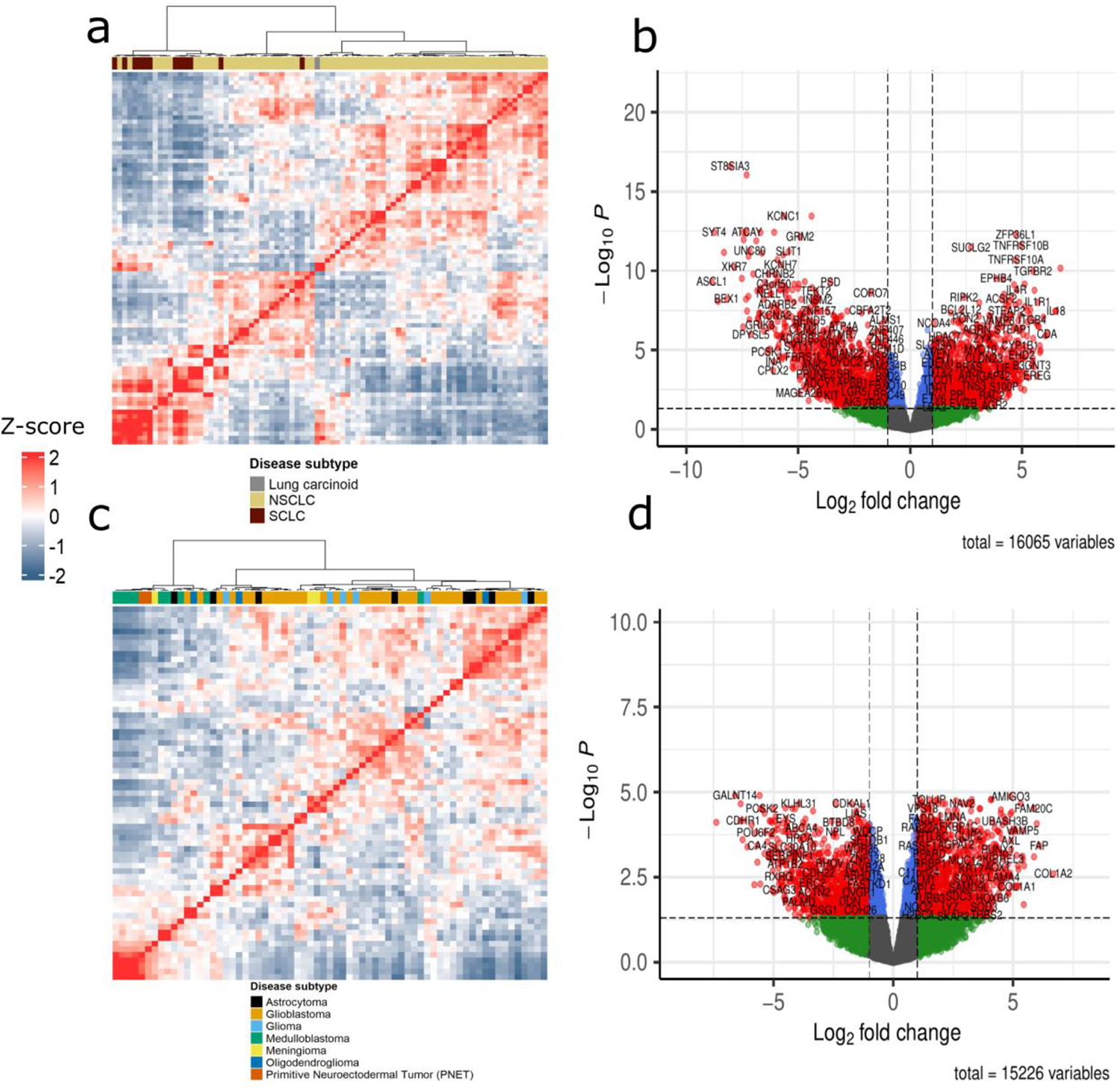
Lung and brain cancer cell lines exhibit extensive transcriptional differences between subtypes. a) Heatmap showing hierarchical clustering of Z-scores of pairwise Spearman correlations of gene expression in lung samples using Ward’s linkage. **b)** Volcano plot of DE genes in NSCLC versus SCLC. Significantly DE genes are colored in red (p-adj <= 0.05 & |LFC| > 1). **c)** Heatmap showing hierarchical clustering of Z-scores of pairwise Spearman correlations of gene expression in brain samples using Ward’s linkage. **d)** Volcano plot of DE genes in glioblastoma versus medulloblastoma. Significantly differentially expressed genes are colored in red. (DE: differentially expressed; NSCLC: non-small cell lung carcinoma; SCLC: small-cell lung carcinoma; p-adj: adjusted p-value; LFC: log fold change).

### Different single-sample network inference methods generate distinct network topologies

First, we aimed to explore the network topology of the aggregate and single-sample networks^34^. **Suppl. figure 1** shows the distribution of edge weights in the aggregate networks as well as across all edges in single-sample networks. Also, it shows the distribution of the average weight of each edge individually across all samples. For lung samples, edge weights in SSN networks ranged between [-0.3, 0.35], while edge weights in LIONESS networks ranged between [-25, 30]. iENA lung networks have an edge weight distribution similar to those constructed by LIONESS, with weights ranging between [-25, 32]. CSN produces networks with binary weights of either zero or one, such that all edges present in the network of a specific sample have a weight of exactly one. Finally, networks constructed by SSPGI had edge weights in the interval [-15000, 15000], with non-continuous values since it is a rank-based method. We observed similar edge weight distributions in networks constructed for brain samples. In both tissues, networks constructed by SSN, LIONESS and iENA were predominantly characterized by edge weights close to zero. Due to the binary nature of edge weights in CSN networks, either zero or one, a significant proportion of edges within each single-sample CSN network was associated with weight zero and thus absent in the network. On average, these networks contained 27 814 and 24 399 edges for lung and brain samples, respectively. We therefore selected the top 25 000 edges in SSN, LIONESS, iENA and SSPGI networks, rendering networks comparable in size. These networks, which had previously already been pruned for HumanNet edges, are further referred to as top 25k networks.

**Figure 2.**
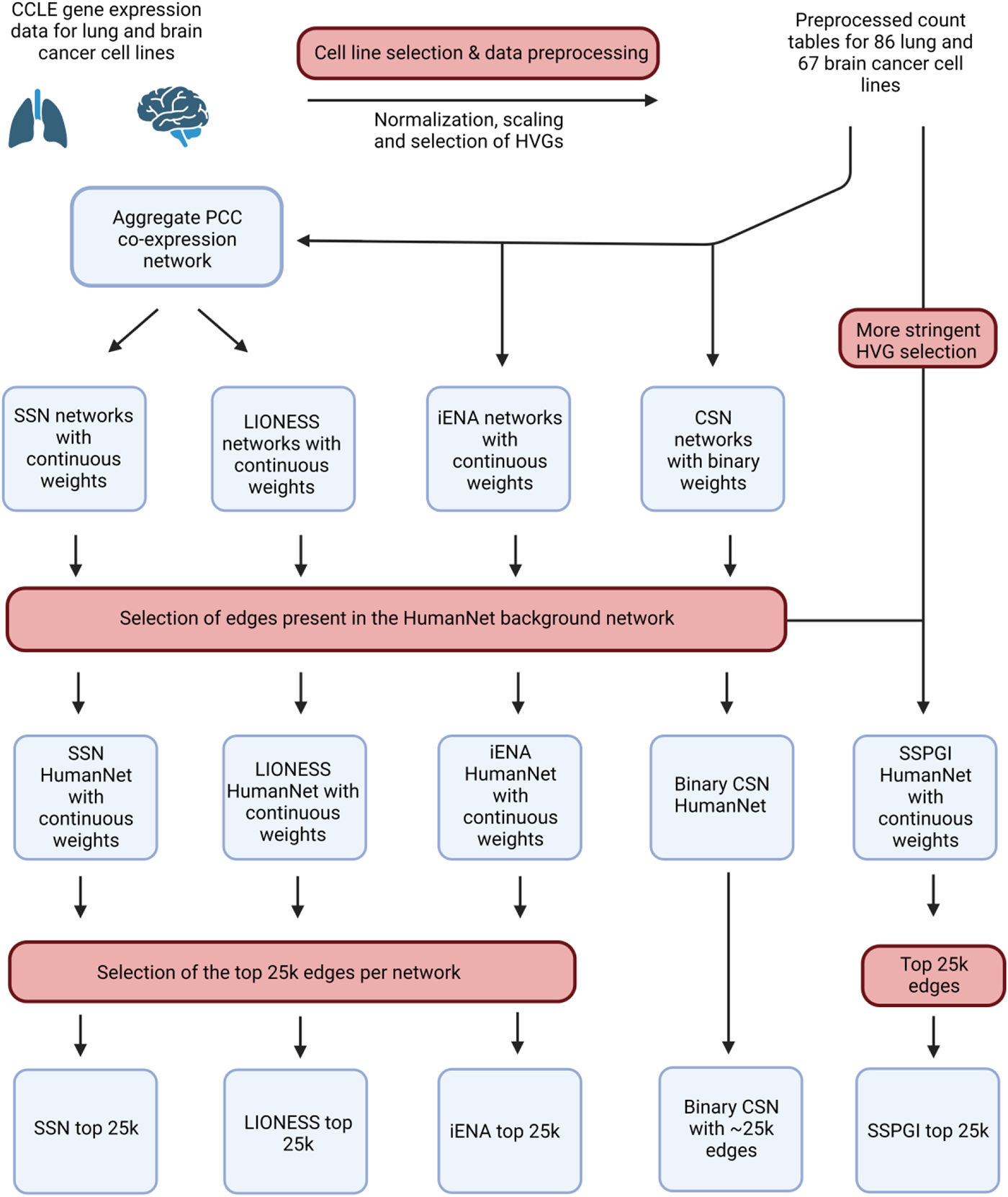
Overview of single-sample network construction and network pruning. Expression data of lung and brain cancer cell lines were downloaded from CCLE, after which samples were selected and data preprocessed (methods). An aggregate co-expression network was constructed for both tissues using Pearson’s correlation (PCC), and single-sample networks using SSN, LIONESS, iENA, CSN and SSPGI. Single-sample networks were pruned for edges present in the HumanNet network and the top 25k edges were selected in SSN, LIONESS, iENA and SSPGI networks, whereas all ∼25k edges were selected in CSN networks. (HVG: highly-variable gene).

The top 25k networks varied in edge weight distributions as well as network topology (**Suppl. Table 1–2**). The aggregate networks had an order of magnitude more connected components than single-sample networks, and also displayed higher clustering coefficients and lower node and edge betweenness. Thus, aggregate networks are more tightly connected with shorter paths between nodes. Overall, topological differences between single-sample networks themselves were rather small, with SPPGI having a lower clustering coefficient than the rest for both tumor types. Thus, although constructed on the same data as the aggregate networks and subjected to similar edge selection procedures, each single-sample network inference method built distinct single-sample networks that are different from the aggregate network, both at the level of edge weight distribution and network topology.

### Clustering of single-sample networks

Next, we inquired to what extent LIONESS, SSN, iENA, CSN and SSPGI provide relevant biological insights at the sample-specific and subtype-specific level. Therefore, we first projected edge weights of single-sample networks onto their first two principal components using Principle Component Analysis (PCA). For lung top 25k networks, NSCLC samples clustered tightly together when single-sample networks were constructed using LIONESS, SSN, or iENA, while SSPGI and CSN networks produced a cloud-like pattern comprising all subtypes (**Figure 3**). Similarly, when projecting edge weights of the top 25k brain networks onto their first two PCs, we observed a dense cluster of astrocytoma, glioblastoma, glioma, meningioma and oligodendroglioma samples next to several medulloblastoma samples for networks constructed using LIONESS, SSN and iENA, while networks constructed using SSPGI and CSN resulted in a cloud-like pattern (**Suppl. figure 2**). The total amount of variance captured by the first two PCs was maximally 25% for both lung and brain top 25k networks, except for iENA brain. To get a more quantitative view of clustering performance based on PCA, we calculated the adjusted Rand index between true subtype classification and a classification after k-means clustering on PC loadings (a score of one indicates a perfect classification) (**Table 1**). We choose to only retain the predominant subtypes glioblastoma and medulloblastoma for brain networks to make the analysis more comprehensible. Adjusted Rand indices were identical between SSN and LIONESS, for both tissues. IENA top 25k networks returned the highest Rand indices, followed by SSN and LIONESS, suggesting that iENA achieves the best separation of samples into disease subtypes. The Rand indices of CSN and SSPGI are notably lower. Thus, this exploratory analysis suggests that some single-sample networks, mainly those constructed with SSN, LIONESS and iENA, reflect more closely the respective disease subtype of a given sample.

**Table 1.** Adjusted Rand indices of k-means clustering on principal components of expression data and the edge weights in single sample networks. Samples were clustered using k-means clustering on PC loadings of the expression data or sample-specific networks, after which cluster allocation was compared to true sample subtypes. Only glioblastoma and medulloblastoma samples were retained for brain networks (PC: principal component).

|  | Expression | SSN | LIONESS | iENA | CSN | SSPGI |
| --- | --- | --- | --- | --- | --- | --- |
| **Lung** | 0.098 | 0.627 | 0.627 | 0.644 | 0.161 | 0.126 |
| **Brain** | 0.092 | 0.363 | 0.363 | 0.475 | 0.189 | 0.069 |

**Figure 3.**
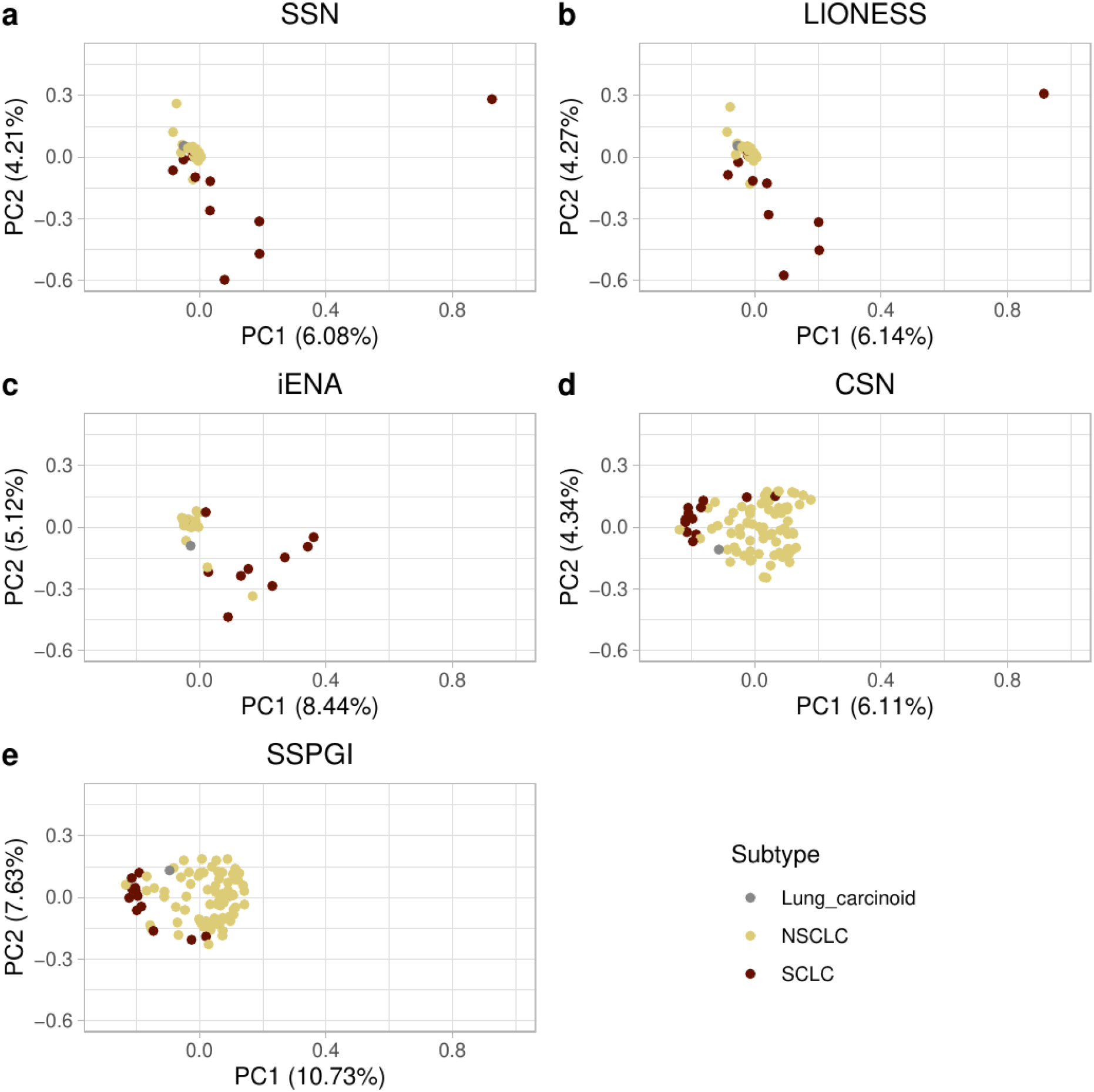
Visualization of lung samples after projecting edge weights onto their first two principal components. We performed PCA analysis on top 25k lung networks constructed using SSN (**a**), LIONESS (**b**), iENA (**c**), CSN (**d**) and SSPGI (**e**). Each dot represents one single-sample network constructed from a cell line corresponding to a given cancer subtype. (NSCLC = non-small cell lung carcinoma, SCLC = small cell lung carcinoma; PCA: principal component analysis).

### Analysis of hubs in single-sample networks

We further identified hubs by selecting the top 200 most connected nodes in each single-sample network and aggregate networks (Methods). As single-sample network inference methods are designed to capture heterogeneity between samples of a tumor type, we expect to some extent different hubs in different samples, and ideally these hub genes are related to the cancer subtype of a given sample. To test this, we first assessed the recurrence of hub genes (i.e. the number of times a given gene is identified as hub across a group of samples) across lung samples in networks constructed using a given method (**Figure 4A**). All methods construct single-sample networks with the majority of hubs being unique to only one or a few samples. However, CSN and SSPGI have 96 and 18 hub genes respectively recurring in all samples. Some hub genes are regularly recurring within SSN, LIONESS and iENA networks, but none overlap across all samples. Furthermore, the top 200 hub genes of the lung aggregate network are consistently recurring among the hub gene sets identified in single-sample networks. Similar observations were made in networks constructed for brain samples, with 76 and 21 hubs overlapping across all samples in CSN and SSPGI networks, respectively, and zero in networks constructed by SSN, LIONESS and iENA (**Figure 4B**). Hubs identified in the aggregate brain network again had a higher incidence of recurrence among hubs identified in single-sample networks. Together, these observations suggest that SSN, LIONESS and iENA produce networks that are inherently more different from each other than CSN or SSPGI networks. Also, hubs in the aggregate network tend to be hubs in single-sample networks.

Hub genes should ideally be related to the cancer subtype of a given sample, and thus similar hubs should be found within sample groups. We thus grouped all NSCLC, SCLC, glioblastoma and medulloblastoma samples and assessed the hub overlap within these groups (**Suppl. Table 3**). Surprisingly, in both the SSN and iENA networks, there were zero common hubs across all samples of any cancer subtype. However, all single-sample networks do have regularly recurring hubs per cancer subtype group (occurring in at least 75% of the samples in a given group) (**Figure 4C-F**). To investigate whether these regularly recurring hubs were subtype-specific, we compared regularly recurring hubs per sample group to recurring hubs across all samples. SSN networks consistently had the highest proportion of subtype-specific recurring hubs compared to the total number of regularly recurring hubs per sample group, followed by iENA and LIONESS. In Figure 4, each dot represent one gene that was identified as a hub, and the y-axis represents the number of times that given hub is found across a given sample group. The highest proportion of subtype-specific versus non-subtype-specific hubs among highly-recurring hubs is thus observed for SSN networks, followed by iENA and LIONESS. CSN and SSPGI have recurring hub genes that were less specific to the cancer subtype of a given sample group (**Figure 4C-F**). Next, we assessed whether these hub lists were enriched for known cancer driver genes. We downloaded a list of known drivers from IntOGen and Cancer Gene Census for NSCLC, SCLC, glioblastoma and medulloblastoma, and assessed their presence in the sets of hubs per sample group. Out of 69 and 59 driver genes for NSCLC and SCLC respectively, 23 were present in the aggregate HumanNet lung network. Known drivers for medulloblastoma and glioblastoma respectively comprised 57 and 45 genes, of which 9 and 16 respectively were present in the aggregate HumanNet brain network. Overall, we found that most methods constructed networks in which hub genes were enriched for subtype-specific cancer driver genes, after concatenating all hubs identified per sample group (**Figure 5 & Table 3**). There was however no tool that clearly outperformed the others across all four cancer subtypes.

**Figure 4.**
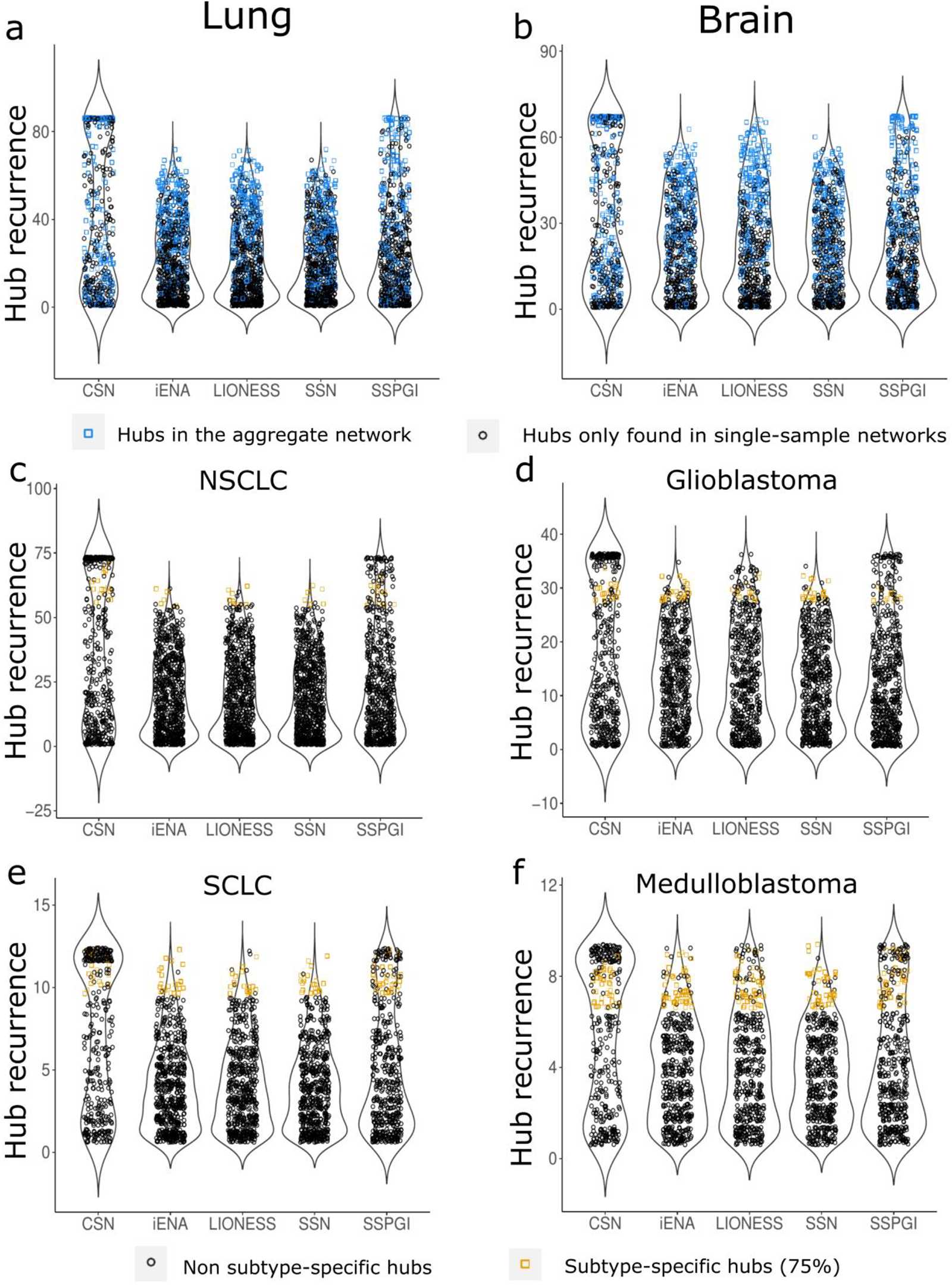
Hub recurrence in single-sample networks. Each dot represents one gene identified as hub, and the y-axis shows how many times a hub recurs in a sample group. **a-b)** Plots containing all hubs over all single-sample networks per inference method for lung and brain respectively. Hubs agreeing to the 200 hubs retrieved from the aggregate network of the same tissue are colored in blue. **c,d,e,f)** Plots containing all hubs over all single-sample networks of one subtype per inference method for lung NSCLC (non-small cell lung cancer) and SCLC (small cell lung cancer), brain glioblastoma and medulloblastoma respectively. For each subtype, the subtype-specific hubs are colored in yellow and defined as the hubs (i) occurring in at least 75% of the networks of that subtype and (ii) the hubs selected in (i) that were not overlapping with the selected hubs of the other subtype for the same tissue.

**Figure 5.**
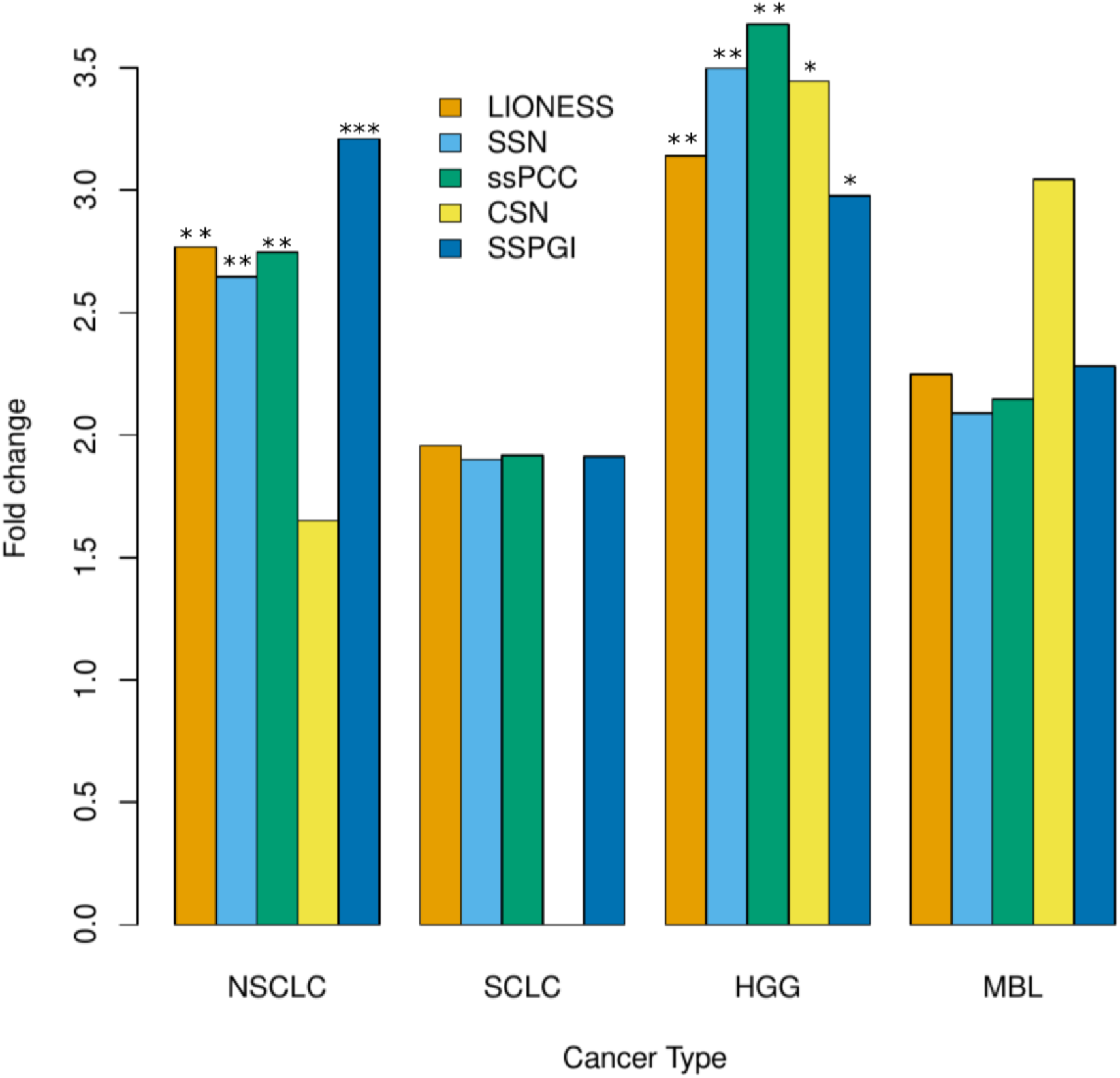
Enrichment of known subtype-specific cancer driver genes in hub gene sets. The top200 most connected nodes in each single sample network were identified as hub genes. Hubs genes were then grouped per sample type and enrichment for known subtype specific cancer driver genes was assessed. (**: p < 0.01, ***: p < 0.001, NSCLC: non-small cell lung cancer, SCLC: small cell lung cancer, HGG: glioblastoma, MBL: medulloblastoma).

We finally assessed whether the same genes were identified as hubs for a given sample across networks constructed by different methods. Networks for lung samples inferred using iENA had, on average, the highest number of hub genes that were not identified as hubs in other methods. In contrast, for LIONESS and SSN, the majority of hub genes were also identified by at least one other method (**Suppl. figure 3A**). Hubs identified by both LIONESS and SSN represented, on average, the largest overlapping group, suggesting again that SSN and LIONESS single-sample networks were most similar. An average of about 35 hub genes were overlapping between all five methods, indicating that all methods prioritized at least a common subgroup of nodes in their respective networks for a given sample. For brain samples, iENA networks again had the largest number of hubs not overlapping with those identified in networks constructed by other methods. Surprisingly, the largest overlap was observed across all five methods, with on average 43 overlapping hub genes across the five networks constructed for a given sample (**Suppl. figure 3B**). Taken together, for a given sample, different methods constructed single-sample networks with different hubs, which also differed from those identified in the aggregate networks.

**Table 2:** Overlap between hubs identified per sample group and known subtype-specific driver genes. The top 200 most connected nodes in each single-sample network were selected as hub genes. These were then grouped per sample group (SCLC, NSCLC, medulloblastoma or glioblastoma), and the intersection with known subtype-specific driver genes was determined.

|  | **SSN** | **LIONESS** | **iENA** | **CSN** | **SSPGI** |
| --- | --- | --- | --- | --- | --- |
| **SCLC** | ERB4, EGFR, SOX2, NOTCH1, FGFR2 | ERB4, EGFR, SOX2, NOTCH1, FGFR2 | ERB4, EGFR, SOX2, NOTCH1, FGFR2 | / | EGFR, NOTCH1, PTPN13, FGFR2 |
| **NSCLC** | ERB4, EGFR, SLC34A2, SOX2, NOTCH1, KDR, PTPN13, CDKN2A, FGFR2 | ERB4, EGFR, SLC34A2, SOX2, NOTCH1, KDR, PTPN13, CDKN2A, FGFR2 | ERB4, EGFR, SLC34A2, SOX2, NOTCH1, KDR, PTPN13, CDKN2A, FGFR2 | EGFR, NOTCH1, FGFR2 | ERB4, EGFR, SLC34A2, SOX2, NOTCH1, KDR, PTPN13, CDKN2A, FGFR2 |
| **Glioblastoma** | CACNA1D, EGFR, PDGFRA, FGFR4, CDKN2A, COL1A1, TP53 | CACNA1D, EGFR, PDGFRA, FGFR4, COL1A1, TP53 | CACNA1D, EGFR, PDGFRA, FGFR4, CDKN2A, COL1A1, TP53 | / | CACNA1D, EGFR, PDGFRA, FGFR4, COL1A1, TP53 |
| **Medulloblastoma** | WNK4, TP53 | WNK4, TP53 | WNK4, TP53 | WNK4, TP53 | WNK4, TP53 |

### Differential node importance in single-sample networks

Due to the use of PCC as underlying network inference method, all single-sample networks were undirected. Thus, instead of evaluating differential targeting ^35^, we defined the ‘node importance’ of a given node as the sum of absolute edge weights of that node, after scaling weights to values between -1 and 1. Using linear modeling and an empirical Bayes procedure, we identified 59 differentially important nodes (p-adj < 0.05 & |LFC| > 1) between SCLC and NSCLC samples in LIONESS networks, 192 in SSN networks and 363 in SSPGI networks. Only one node was significantly differentially important in CSN lung networks, and none in iENA networks (**Figure 6**). However, none of these gene sets were enriched for NSCLC- and SCLC-specific known driver genes. For brain networks, we found 113, 116, and 178 differentially important nodes between glioblastoma and medulloblastoma samples in SSN, LIONESS and SSPGI networks respectively (**Suppl. figure 4**). Here we observed a significant enrichment for known medulloblastoma and glioblastoma driver genes in SSPGI networks, but not in others. There was a strong tendency towards negative LFCs in SSN and LIONESS networks, a phenomenon not observed during DE analysis. This observed preference is likely caused by an unbalanced group size of samples used to construct the aggregate network, i.e. 73 NSCLC versus 12 SCLC samples and 53 glioblastoma versus 9 medulloblastoma samples, resulting in aggregate networks which are more representative of NSCLC and glioblastoma samples respectively (**Suppl. Figures 5–14**).

**Figure 6.**
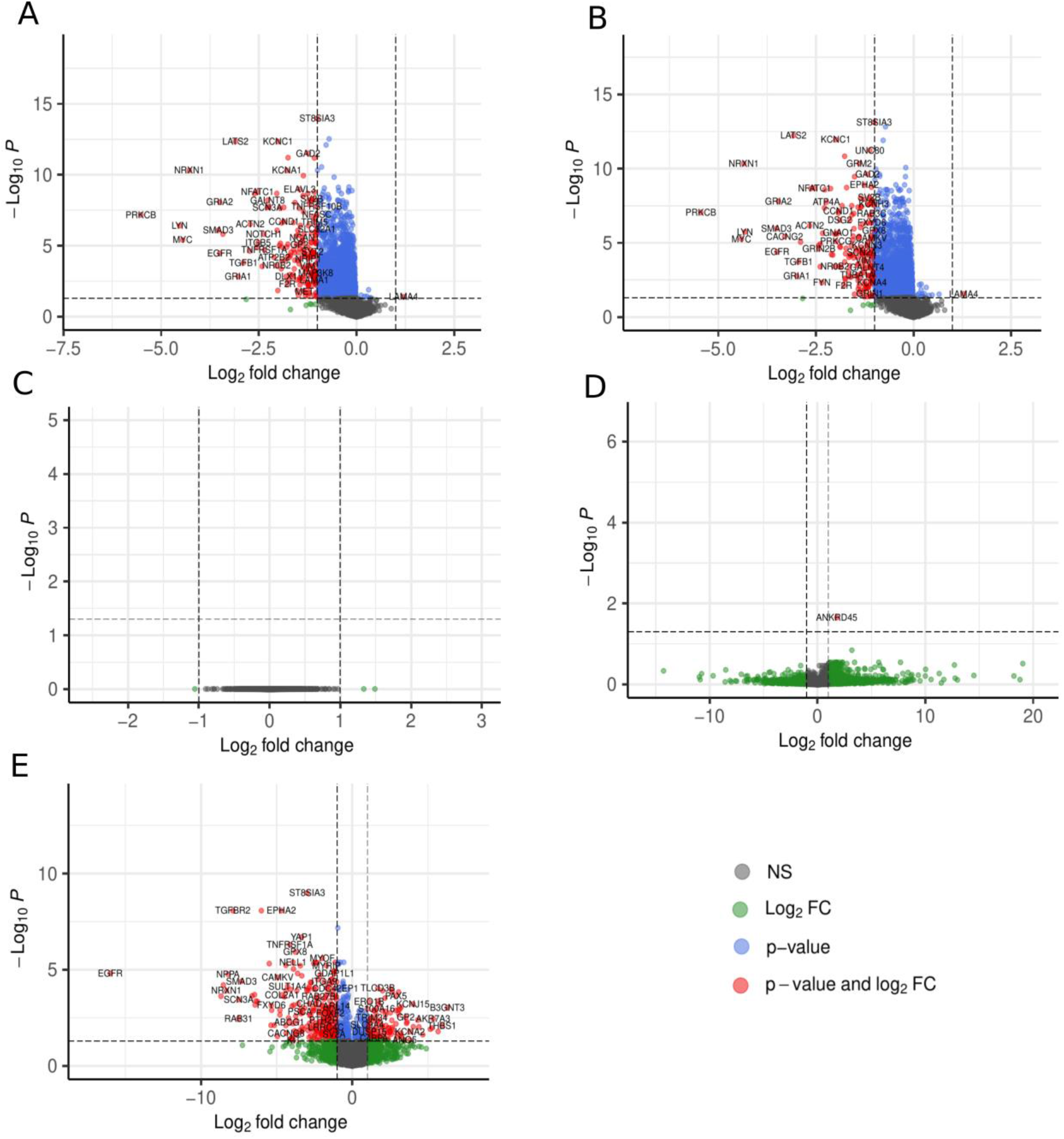
Single-sample networks display distinct differential node importance across network types. The node importance, or sum of absolute edge weights, was calculated for all nodes in top 25k networks constructed by SSN (A), LIONESS (B), iENA (C), CSN (D) and SSPG (E)I. Differentially important nodes (p-adj < 0.05 & |LFC| >= 1) in small cell lung carcinoma versus non-small cell lung carcinoma were identified using linear modelling and an empirical Bayes procedure. (p-adj: adjusted p-value; LFC: log fold change, NS: non-significant).

### Relating single-sample networks to sample-specific molecular features

Finally, we assessed the biological relevance of single-sample networks by comparing them to additional CCLE omics measured on the same samples. Ideally, these single-sample networks constructed from transcriptional gene expression profiles have a higher resemblance to other sample-specific omics than the aggregate network has. We downloaded proteomics and copy number variation (CNV) data from CCLE (Methods) and assessed the correlation between node importance and protein abundance, and node importance and CNV (**Figure 7A-D**). For the aggregate networks, we assessed correlations between node importance in the aggregate network and proteomics/CNV measurements in individual samples, while for single-sample networks we calculated node importance in each network separately. On average, node importance in the aggregate network does not correlate well with protein abundance or CNV data, displaying correlation coefficients < 0.1 for proteomics data and < 0.05 for CNV data. On the other hand, for all methods, single-sample networks outperformed the aggregate network in respect to correlation of node importance to both proteomics or CNV data. Only for brain single-sample networks constructed by CSN we detected no significant increase in the average correlation coefficient between the sum of absolute edge weights and protein abundance. Overall, single-sample networks from SSN and LIONESS resulted in the biggest increase in average correlation coefficients, for both lung and brain samples, and for proteomics and copy number variation data. Together, these findings suggest that single-sample network inference methods were better in capturing sample-specific molecular features than aggregate networks and that among the applied single-sample network inference methods, SSN and LIONESS are most capable of capturing sample-specific protein abundance and CNV features.

**Figure 7.**
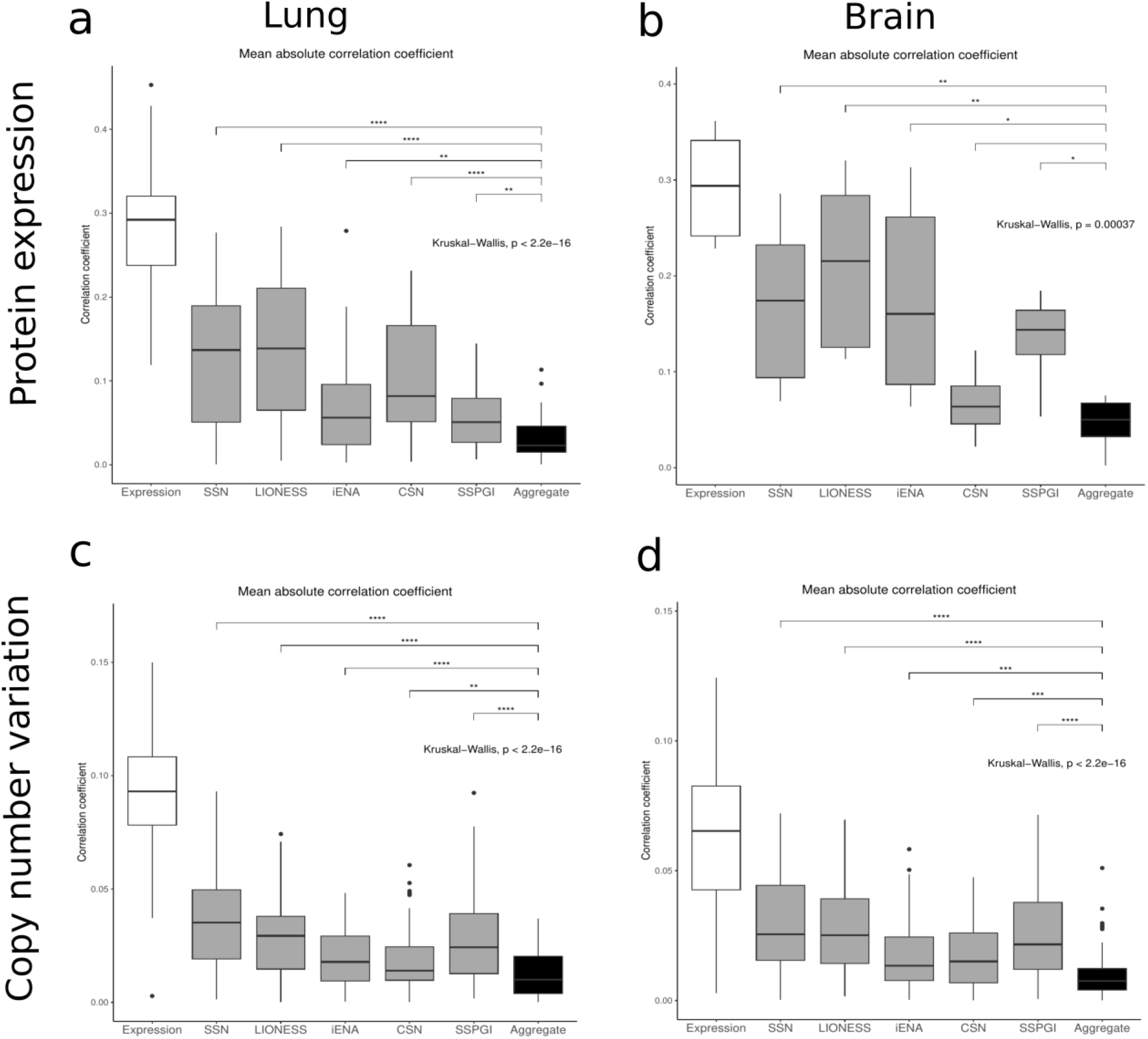
Feature-wise correlation between single-sample networks and other omics data. **a)** Lung proteomics data; **b)** brain proteomics data; **c)** lung Copy Number Variation (CNV) data; **d)** brain Copy Number Variation data. Differences of the average correlation coefficients between the single-sample methods and the aggregate network is shown above the boxplots (* p ≤0.05, ** p ≤ 0.01, *** p ≤ 0.001 and **** p≤ 0.0001, Kruskal-Wallis test).

## Discussion

The fight against highly complex and heterogeneous diseases such as cancer necessitates an in-depth understanding of disease pathobiology, at population level, but especially at the level of individual patients. Investigation of gene networks and their rewiring in disease can therefore greatly benefit the development of individualized therapeutic strategies. Although single cell technologies offer the ability of constructing gene networks for individual patients, there are still limitations associated with this approach, especially in the clinical setting. Bulk molecular profiling techniques on the other hand are well established, are cheaper, there is a plethora of data already available and network inference algorithms have been extensively benchmarked^12^. However, bulk network inference methods construct population-level networks, representing interactions shared by most patients. Single-sample network inference methods have thus been developed to prioritize biologically meaningful information of a single individual, bridging the gap towards personalized medicine, a major goal in present-day cancer research^36^.

In this study, we compared five single-sample network inference algorithms, LIONESS, SSN, iENA, CSN and SSPGI in their construction of single-sample networks, with regard to graph properties, biological relevance of hubs, the ability to distinguish samples of different cancer subtypes from each other, as well as their concordance with other sample-specific omics data. Although each method has been shown to function as intended in its original publication and research context, these studies understandably lack neutrality, and so far only limited benchmarking/comparison has been performed. First, we discovered that each method had different characteristics and requirements concerning in- and output data structures. The SSPGI algorithm was not scalable above 7800 genes. Also, as CSN returns binary networks comprising either zero or one as edge weights, the average number of edges within each single-sample network was lower compared to other methods. Therefore, we first pruned networks by selecting for edges present in the HumanNet reference network, and then selected the top 25 000 edges within each single-sample network. Furthermore, edge weights ranged between highly different outer bounds in networks constructed by SSN and SSPGI, so we scaled them to values between [-1,1] for all methods.

Existing benchmarks focused on a limited number of features per sample, on the performance of further downstream structural control (SSC) methods, or only evaluated a limited number of network inference tools^27–29^. A comparison between LIONESS and SSN revealed that when both methods depend on the exact same aggregate network, there is an almost perfect linear relationship between edge weights of LIONESS and SSN networks for a given sample^29^. Both methods heavily rely on PCC for network construction, and construct highly similar networks. Indeed, we found that SSN and LIONESS networks had similar network characteristics, hub gene sets and classification performances.

One critical remark is that the original applications of LIONESS, SSN, iENA and SSPGI used a group of healthy samples to create the aggregate network, which was not the case in this study. Paired healthy and disease samples are not always available in a clinical setting and not for all tumor types, thus we aimed to investigate the performance of these methods in the absence of control reference samples. When the aggregate network is constructed from a healthy or homogenous group of samples, each sample of interest will be compared to this aggregate network representing a healthy state. One can thus argue that the construction of an aggregate network from a heterogenous group of samples will eventually result in less explicit differences between the aggregate and the single-sample networks. This is partly reflected in our differential node importance analysis. Due to the unbalanced sample numbers during the construction of aggregate lung (73 NSCLC versus 12 SCLC samples) and brain (53 glioblastoma versus 9 medulloblastoma samples) networks, the final aggregate networks are dominated by the larger sample group. As a result, we observed higher average absolute edge weights for samples belonging to the underrepresented group. However, the effects of using a homogenous or heterogenous group of reference samples on classification performance has previously been investigated and was found to be only minor^29^.

Yet, we advocate for the careful assembly of the aggregate networks with subgroups of similar size. Ideally, also covariates such as cell culture type, age, gender… are taken into account^37^. Hub genes in SSN networks, identified based on node degree, have been shown to be strongly related to cancer driver mutations^19^. However, there is no consensus regarding the number of nodes to select as hubs. While the SSN study suggests to use the top 5, 10 or 20 most connected nodes, we selected the top 200 most connected nodes and found hub gene sets to be significantly enriched for known subtype-specific driver genes. Within methods, there was a variable number of overlapping hubs between different single-sample networks and the aggregate network, with hub genes identified in CSN networks displaying the lowest diversity across samples. As a result, CSN hubs also had the lowest cancer subtype specificity, as most hubs regularly recurred across both subtypes. Since all single-sample networks were undirected coexpression networks, we identified node importance as the sum of absolute edge weights. The number of differentially important nodes in single-sample networks differed highly between methods, although these were not enriched for known subtype-specific cancer driver genes.

Overall, there is a lack of ground truth data which makes a true benchmark study difficult. Instead, we explored the relationship between single-sample networks and other omics data modalities, namely proteomics and CNV, at the sample level. We found that, in general, single-sample networks inferred by any method, outperformed the aggregate network regarding correlation to other omics. Also, we heavily relied on cancer subtype annotations of CCLE cell lines during our hub gene and differential importance analyses. Yet, clustering of samples based on expression data showed that these annotations might not be ideal, as several samples clustered together with other subtypes. These issues have previously been addressed^32^, and we used a similar approach in combination a clustering analysis (methods) with to select the cell lines/samples included in our study.

In conclusion, we have constructed single-sample networks for 86 lung and 69 brain cancer cell lines from CCLE, using five different single-sample network inference methods. Several network pruning steps were required to make networks comparable. We found that hub genes were enriched for known cancer subtype-specific driver genes, suggesting that single-sample networks are a valuable tool for personalized medicine. Overall, SSN and LIONESS performed consistently across analyses, and these networks were accurate reflections of other omics measured on the same samples. Moreover, the LIONESS framework furthermore allows the use of any network inference algorithm. Hence, from our study, we conclude that single-sample network inference algorithms are an invaluable resource for personalized medicine and precision oncology.

## Methods

### Data and cell line selection

Expression read counts and metadata were downloaded from the DepMap (Cancer Dependency Map) website (20Q4 version 2 release)^30, 31, 38^. Expression data were available for 84 primary brain cancer and 189 primary lung cancer cell lines. For both tumor types, outlier cell lines were excluded in two ways. First, only cell lines with Spearman correlation of expression profiles greater than 0.55 to real tumor tissue, as outlined in a pan-cancer comparison of CCLE cell lines and TCGA tumor samples^32^, were kept to ensure biological interpretability. Second, cell lines strongly differing from the other cell lines of the same tumor type were removed by clustering to exclude unrepresentative samples for a given tissue. Clustering was done with the hclust function using average agglomeration, with tree cutting at 95% of the maximum height of the tree in R (version 3.6). Clusters with less than 3 samples were removed. After filtering, 67 brain and 86 lung cancer cell lines were retained for single-sample network inference.

### Expression data preprocessing

The RNA-seq count data was processed using EdgeR^39^ in R (version 3.6). Raw counts were filtered to keep only genes with counts per million (cpm) greater than 1 in at least one sample. Next, raw counts were normalized by library size, converted into cpm and log-transformed. We selected 7942 and 9252 highly variable genes (HVG; variance >2.75 over all samples per tumor type), respectively, for brain and lung as input for single-sample network inference (HVG selection). Due to lack of scalability, only 7800 highly variable genes were retained for SSPGI for both tumor types (see SSPGI method section). Finally, scaling and centering were performed per gene. Heatmaps were constructed after calculating Spearman correlations between samples and applying Ward linkage using ComplexHeatmap^40^.

### Aggregate networks and network visualization

Aggregate networks were constructed separately for brain and lung samples using PCC after HVG selection. These fully connected networks were subjected to pruning for edges in the HumanNet background network, an integrated functional gene network^33^. We choose to work with the HumanNet-XN v2 network, which contains 17 929 genes and 525 537 edges representing physical protein-protein interactions, functional annotations substantiated by different omics data and interologs from other species and co-citation links. We then selected the top 25 000 edges based on edge weights in these networks, and visualized them in Cytoscape. A list of driver genes for NSCLC, SCLC, glioblastoma and medulloblastoma was downloaded from the IntoGen website (19/06/2023, https://www.intogen.org/search) and the

Cancer Gene Census website (19/06/2023, https://cancer.sanger.ac.uk/cosmic/census), and nodes within a distance of 1 from these known drivers were selected for visualization. The respective nodes were annotated as driver or non-driver genes. We then analyzed the networks using CytoScape’s built-in network analysis functions, and driver genes were colored based on degree.

### Single-sample network inference methods

**Table 3** provides an overview of the different single-sample network inference methods used in this study. We slightly modified several methods to run them with PCC as underlying network inference method, without ‘normal tissue’ reference samples, as well as with different or without background networks.

**Table 3.** Overview of the different single-sample network inference methods used in this study. We slightly modified several methods because we ran them with PCC as underlying network inference method for comparison, without ‘normal tissue’ reference samples or with different or without background networks.

| Method | Underlying concept | Remarks |
| --- | --- | --- |
| **SSN<sup>19</sup>** | Differential correlation between reference network and reference network + sample of interest<br>$\Delta PCC = PCC_{r+1} - PCC_r$<br>with $PCC(x_i, x_j) = \frac{Covariance(x_i, x_j)}{\sqrt{Variance(x_i)Variance(x_j)}}$ | All cancer samples of specific tumor type as reference, no normal samples<br>No background network such as STRING, no significance testing of the edges |
| **LIONESS**<sup>22</sup> | Linear interpolation<br>$e_{i,j}^q = N(e_{i,j}^\alpha - e_{i,j}^{\alpha-q}) + e_{i,j}^{\alpha-q}$<br>With e edge weight in respectively all samples network ($\alpha$) or all samples network without sample of interest q ($\alpha - q$) and N scaling factor inversely proportional to the total number of samples | PCC as aggregate network inference method |
| **IENA**<sup>24</sup> | Single-sample correlation<br>$sPCC(x_i, x_j) = \frac{Covariance^r(x_i, x_j)}{\sqrt{Variance^r(x_i) Variance^r(x_j)}}$<br>with $Covariance^r(x_i, x_j) = (x_i - \mu_i^r)(x_j - \mu_j^r)$<br>with $Variance^r(x_i) = \frac{1}{N} \sum_{i=1}^N (x_i - \mu_i^r)^2$<br>With mean and variance for genes $x_i$ and $x_j$ taken from the reference $r$ | Only sPCC node-networks from iENA, not shPCC edge-networks<br>All cancer samples of specific tumor type as reference, no normal samples |
| **CSN**<sup>26</sup> | Local gene-gene association based on statistical independency model<br>$\rho_{xy}^s = \frac{n_{xy}^s}{n} - \frac{n_x^s}{n} \cdot \frac{n_y^s}{n}$<br>With $\rho$ statistic of edge x-y in cell / sample s, n is the number of cells / samples with expression of gene x, y or both as plotted in scatter diagrams | Binary output<br>Developed for single cell<br>Scatter diagrams per gene pair considering all cancer samples |
| **SSPGI**<sup>25</sup> | Edge perturbation based on gene expression rank subtraction<br>$\Delta_{e,s} = \delta_{e,s} - \bar{\delta}_e$ $\delta_{e,s} = r_{i,s} - r_{j,s}$<br>With expression rank r, delta rank matrix $\delta$ and benchmark delta rank vector $\bar{\delta}$ for an edge e between genes i and j within a sample s | HumanNet as background network, not Reactome<br>Not scalable, up to 7800 genes<br>Benchmark delta rank vector calculated using all cancer samples of specific tumor type as reference, no normal samples |

In **SSN** (Single-Sample Network), a reference PCC network is first generated based on transcriptome data of several reference samples, usually normal tissue samples. Here we used all selected cell lines for a specific tumor type, with the sample of interest each time omitted as a reference. Then, the same is done for all the reference samples plus the sample of interest to generate the so-called perturbed network. Finally, these two networks are subtracted from each other. The significance of p-values is not considered as we prune all the networks in the same way using HumanNet and 25k HumanNet (**Figure 2**.)^19^. Although there is a SSN Python implementation online, we made our own R implementation for ease of use with the above mentioned modifications.

In **LIONESS**, linear interpolation is performed on the edge weights of two networks, here constructed by PCC, the first containing all samples, the second containing all samples except for the sample of interest^41^. We used the LIONESS function from the LIONESS R package (https://github.com/mararie/LIONESSR). This function creates one edge list file for all samples in the input expression dataset. We used the single-sample PCC calculation in the iENA node-network^24^, where the PCC between two genes in a single-sample is calculated using mean and variance in a reference group, usually normal tissue samples, but here all selected cell lines for a specific tumor type. A customized implementation of the algorithm in R was made because no source code was provided with the original publication.

The **Cell-Specific Network** (CSN) was developed originally to infer gene association networks for single cells. Still, it can also be applied to bulk data to infer single-sample networks^26^. It generates a binary output, where gene-gene interactions are considered present (1) or not present (0). CSN is based on statistical dependency. For each gene pair, an expression scatter diagram is made for which each dot represents one cell or sample. Then three different neighborhoods are defined around the expression of gene x, gene y and the expression of both genes based on predefined distance parameters. The number of neighboring cells or samples in these neighborhoods (nx, ny and nxy) divided by the total number of cells or samples n are estimates of the marginal density function of x and y and the joint density function of x and y, respectively. These are used to define a statistic from -1 to 1, which follows a normal distribution given that gene x and gene y are independent. Because the mean and the standard deviation of this normal distribution are known, this statistic can be used to calculate a p-value that, in case of significance, rejects the null hypothesis that gene x and y are independent in sample k and form an edge. The MATLAB code for this method is provided on the papers GitHub page (https://github.com/wys8c764/CSN).

**SSPGI** calculates edge perturbation values^42^. First, the gene expression matrix is converted into a rank matrix by ranking the genes according to the expression value in each sample. Second, a delta rank matrix is calculated by subtracting the ranks of any two genes connected by a given edge in the background network. The original publication created a background network based on all gene interactions in the Reactome pathway database^43^. In theory, the required background network could contain all possible edges between all genes in all samples, but in practice this is not feasible due to the lack of scalability of SSPGI. We could run SSPGI only on 7800 genes and with HumanNet given as background network, which causes no other interactions being calculated than the ones present int HumanNet. Including more genes, or all possible edges as background caused the method to terminate with an error. For all methods we worked on a high performance computing infrastructure on an Dual Intel Xeon Gold 6420 CPU cluster using one node with a usable memory of 700 GBM and 2x18 cores. As within-sample delta ranks of gene pairs are stable under normal conditions, a benchmark delta rank vector is calculated using the mean rank of all genes across the reference group of normal samples. However, we built this benchmark delta rank vector using all selected cell lines for a specific tumor type. Finally, the edge perturbation matrix is created by subtracting every samples benchmark delta rank vector from the delta rank matrix. The authors provide the SSPGI implementation written in R on their GitHub page (https://github.com/Marscolono/SSPGI).

Outputs from each single-sample network inference method were converted into a dataframe with edges as rows and samples as columns, generating a uniform format as provided by the LIONESS algorithm. In a subsequent step, we filtered the edges of the other single-sample networks obtained by SSN, LIONESS, iENA and CSN based on the HumanNet background network, as explained for the aggregate network. In this way, these networks were comparable to those constructed by SSPGI.

### Analysis of network topology

The average edge weight distribution was plotted for each method using ggplot2’s *geom_density* function in R^44^. The average weight of an edge was determined by calculating the average weight of a given edge across all samples, ignoring entries for which the given edge was not present in the single-sample network. For plotting the weight density distribution of all edges, the weights of all non-zero edges were concatenated into a single weight vector. We used the igraph package in R to determine network characteristics^45^. Clustering coefficients were calculated using the *transitivity* function with type *average*, network densities using the *edge_density* function without considering loops. Node betweenness was calculated using the *estimate_betweenness* function while treating the network as an undirected graph. As this is a node-specific characteristic, we calculated the mean value across all nodes within each sample. Edge betweenness was determined using the *betweenness* function with the same parameters as for node betweenness. Finally, the *diameter* and *count_components* functions were used to calculate network diameter and the number of connected components, respectively.

### Principal component analysis and calculation of the adjusted Rand index

Top 25k networks were transposed so rows represented samples and columns represented edges, after which R’s *prcomp* function was applied. Plots were drawn using the *autoplot* function in ggplot2, and dots were colored according to the sample cancer subtype. We applied k-means clustering with k = 3 (lung) or k = 2 (brain) on PC loadings, such that every sample was assigned to a cluster. Due to the limited sample sizes for most brain cancer subtypes, only glioblastoma and medulloblastoma samples were retained. Samples were associated with a true cluster based on their cancer subtype in the metadata file, after which the k-means clustering result was compared to the true subtype clustering using the *adj.rand.index* function from the *fossil* library.

### Hub gene analysis

Hub genes were identified as the top 200 most connected nodes in each top 25k single-sample network. Enrichment for known cancer driver genes was assessed under a hypergeometric distribution using a combined list of all known driver genes per tissue and all genes present in the network as background. Violin plots were made visualizing the recurrence of hubs in all samples as well as all samples within a disease subtype. Regularly recurring hubs were defined as hubs recurring in at least 75% of networks in a sample group per method. Next, we determined the overlap between hub genes identified in an individual sample across different single-sample networks and plotted the different combinations using ggplot2.

### Differential node importance

Since LIONESS, SSN, iENA, CSN and SSPGI produce single-sample networks with different ranges of edge weights, we performed a within-sample normalization to scale weights within [-1, 1]. The differential node importance between sample subgroups was evaluated by calculating the sum of absolute edge weights for each node and applying linear modelling with an Empirical Bayes procedure, as implemented in the limma package^46^. The differential important nodes were identified having an absolute log-fold change (LFC) > 1 and an adjusted p-value < 0.05 (Benjamini & Hochberg correction). Enrichment for known cancer driver genes was assessed by a hypergeometric distribution using a combined list of all known driver genes per tissue and all genes present in the network as background.

### Comparison with other omics data

Normalized proteomics datasets were downloaded from the CCLE website (protein_quant_current_normalized.csv, version 20Q4), and cell lines were matched to samples present in single-sample networks for lung and brain, separately. Rows with duplicate gene symbols were removed. As these data were already normalized, no further preprocessing was applied. Next, we selected samples with matching proteomics data from the preprocessed expression dataset. The sum of absolute edge weights was calculated in all single-sample networks and the aggregate networks as described above. We then calculated PCCs between proteomics measurements and the sum of absolute edge weights for each matching sample, and between the aggregate network sum of absolute edge weights and each individual proteomics sample. Finally, sample-wise PCCs were determined between matching samples in the expression data and proteomics data. Copy number variation data was also downloaded from the CCLE website (20Q4_v2_CCLE_gene_cn.csv). We applied the same procedure to copy number variation data as for proteomics. Results were plotted using ggplot2^44^.

## Data and code availability

All scripts used to construct and analyze the regulatory networks in this study can be found at https://github.com/CBIGR/single_sample_networks. No new data was generated for this study.

## Acknowledgements

This study was funded by an FWO PhD fellowship fundamental research [11N5922N] for JD, a BOF PhD fellowship [BOF20/DOC/285] for JL and a BOF Starting Grant [BOF/STA/201909/030]. The funder played no role in study design, data collection, analysis and interpretation of data, or the writing of this manuscript.

## Author information

JD implemented the single sample network inference methods; JD, BV and JL conducted the data analysis and wrote programming code; JD and BV made all figures and tables; JD, BV and VV wrote the initial draft; JD, BV, JL, KDP and VV edited the final manuscript; VV designed and supervised the study; JD, BV, JL and KDP contributed to the conceptualization of the study. All authors read and approved the final manuscript.

## Supplementary materials

**Suppl. Table 1.** Average network characteristics for single-sample networks of lung samples (n = 86) constructed by different single-sample network inference algorithms. ‘# Edges’ refers to the total number of edges in the networks after selection for highly variable genes and selection for edges present in the HumanNet network, while ‘# Edges retained per sample’ refers to the average number of non-zero edges in CSN networks, guiding us to select the top 25000 edges per sample in other networks as well. See methods section for more details on how each network characteristic was calculated.

|  | AGGREGATE | SSN | LIONESS | IENA | CSN | SSPGI |
| --- | --- | --- | --- | --- | --- | --- |
| # NODES | 5454 | 5454 | 5454 | 5454 | 5454 | 4814 |
| # EDGES | 53296 | 53 296 | 53 296 | 53 296 | 53 296 | 42 193 |
| # EDGES<br>RETAINED PER<br>SAMPLE | 25000 | 25 000 | 25 000 | 25 000 | 27 813 | 25 000 |
| CLUSTERING<br>COEFFICIENT | 0.5984 | 0.28 | 0.29 | 0.30 | 0.35 | 0.17 |
| DENSITY | 0.0028 | 0.0023 | 0.0024 | 0.0027 | 0.0035 | 0.0023 |
| NODE<br>BETWEENNESS | 2059.273 | 7198.60 | 7043.12 | 6469.36 | 5556.04 | 7203.90 |
| EDGE<br>BETWEENNESS | 0.0002 | 0.0007 | 0.0007 | 0.0007 | 0.0007 | 0.0007 |
| DIAMETER | 13.5994 | 10.90 | 11.58 | 10.72 | 13.77 | 11.65 |
| CONNECTED<br>COMPONENTS | 739 | 19.73 | 37.97 | 20.11 | 42.27 | 21.83 |

**Suppl. Table 2.**
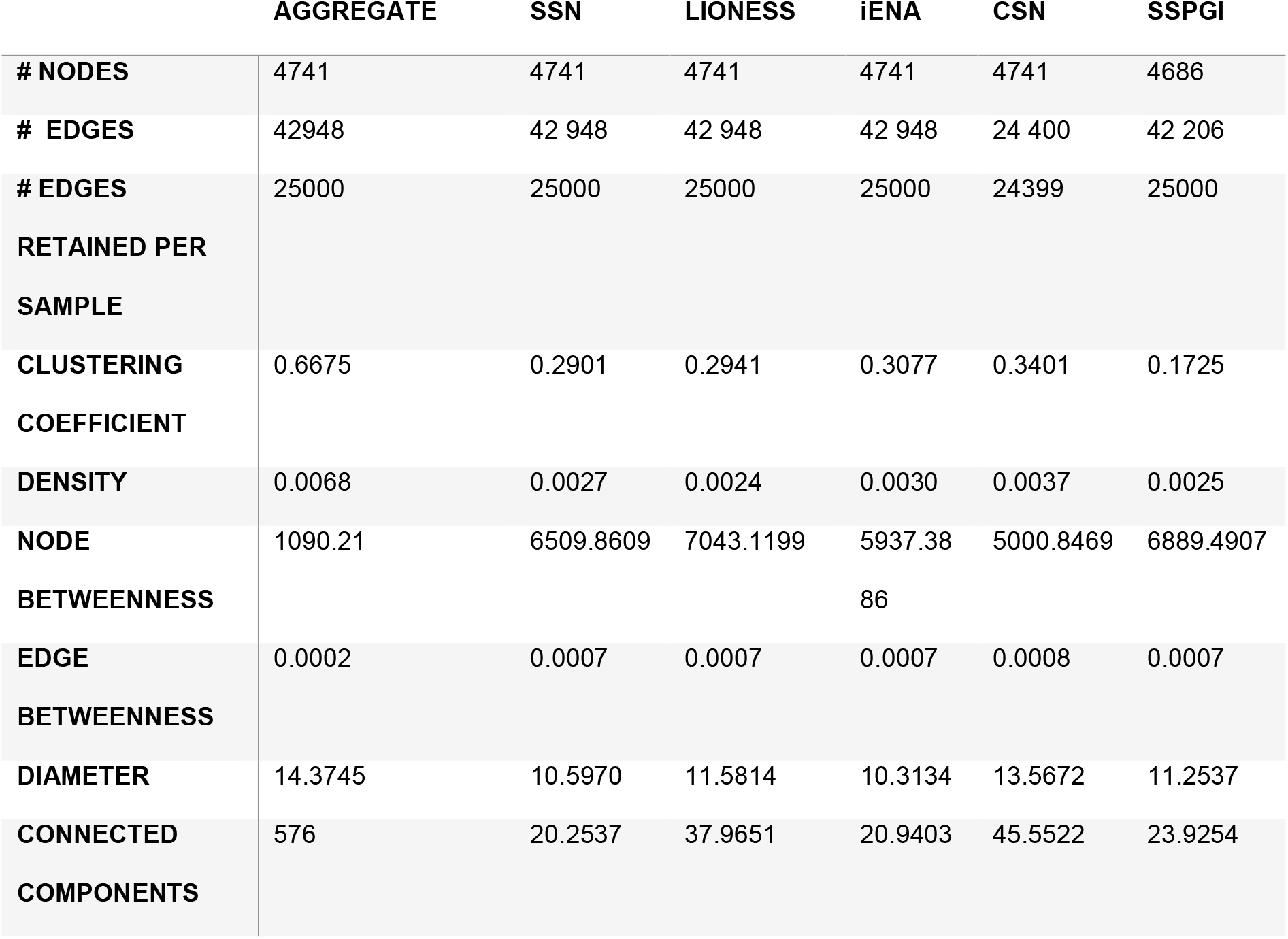
Average network characteristics for single-sample networks of brain samples (n = 67) constructed by different single-sample network inference algorithms. ‘# Edges’ refers to the total number of edges in the networks after selection for highly variable genes and selection for edges present in the HumanNet network, while ‘# Edges retained per sample’ refers to the average number of non-zero edges in CSN networks, guiding us to select the top 25000 edges per sample in other networks as well. See methods section for more details on how each network characteristic was calculated.

**Suppl. Table 3.** Overview of hub recurrence across sample groups and single-sample network inference methods. The top 200 most connected nodes were identified as hubs in each single-sample network, after which the intersection was determined per sample group. (NSCLC = non-small cell lung carcinoma, SCLC = small cell lung carcinoma).

|  | SSN | LIONESS | iENA | CSN | SSPGI |
| --- | --- | --- | --- | --- | --- |
| NSCLC (n=73) | 0 | 0 | 0 | 97 | 18 |
| SCLC (n=12) | 0 | 3 | 0 | 125 | 47 |
| Glioblastoma (n=60) | 0 | 24 | 0 | 100 | 50 |
| Medulloblastoma (n=12) | 0 | 2 | 0 | 85 | 26 |

**Suppl. Figure 1.**
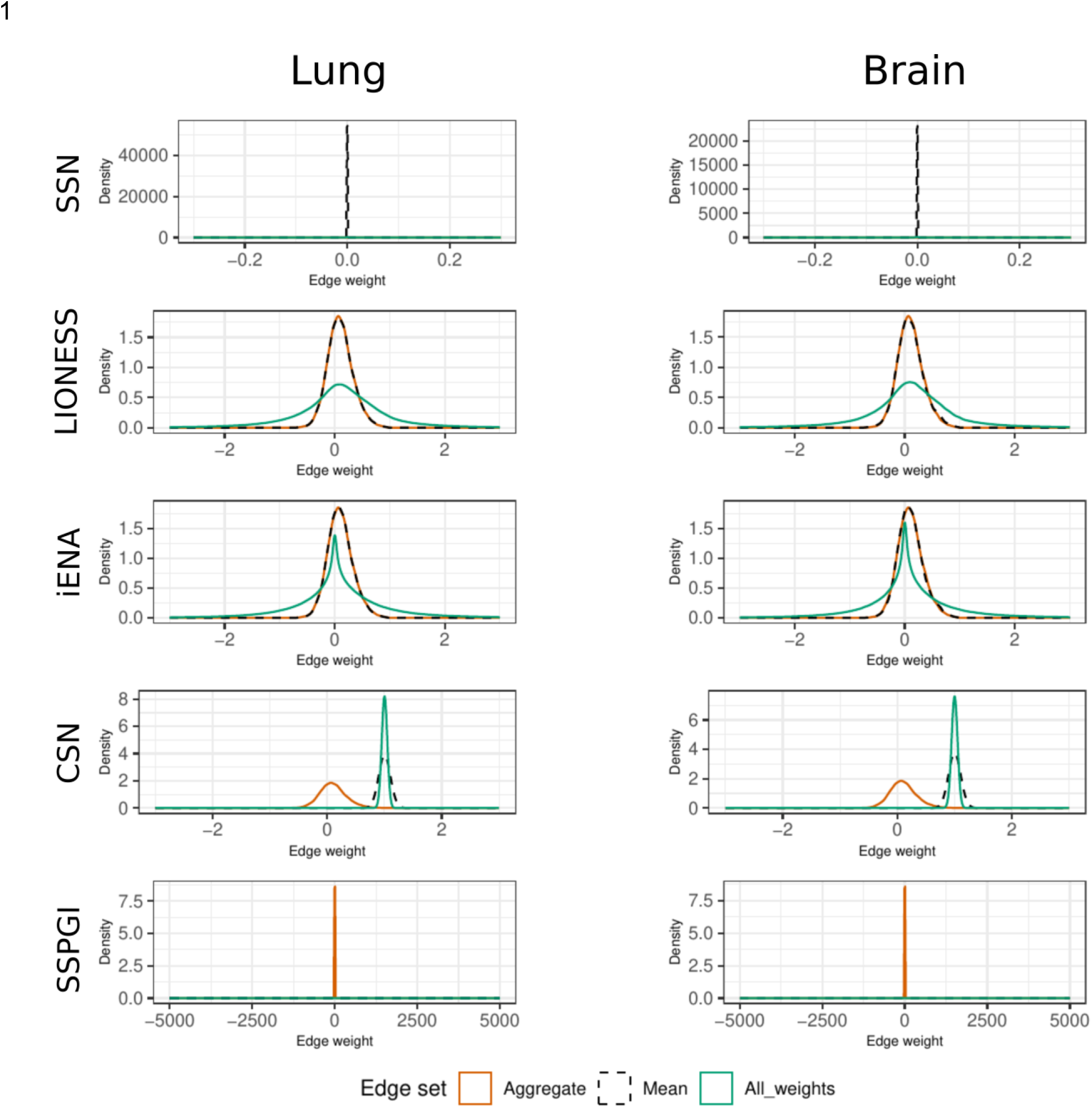
Single-sample networks constructed by SSN, LIONESS, iENA, CSN and SSPGI are characterized by distinct edge weight distributions. Edge weight distributions were plotted after selection for edges present in the HumanNet network, for the aggregate network (orange), the mean edge weight for every single edge (only using non-zero edges) (black) and a concatenation of all weights across every sample (green).

**Suppl. Figure 2.**
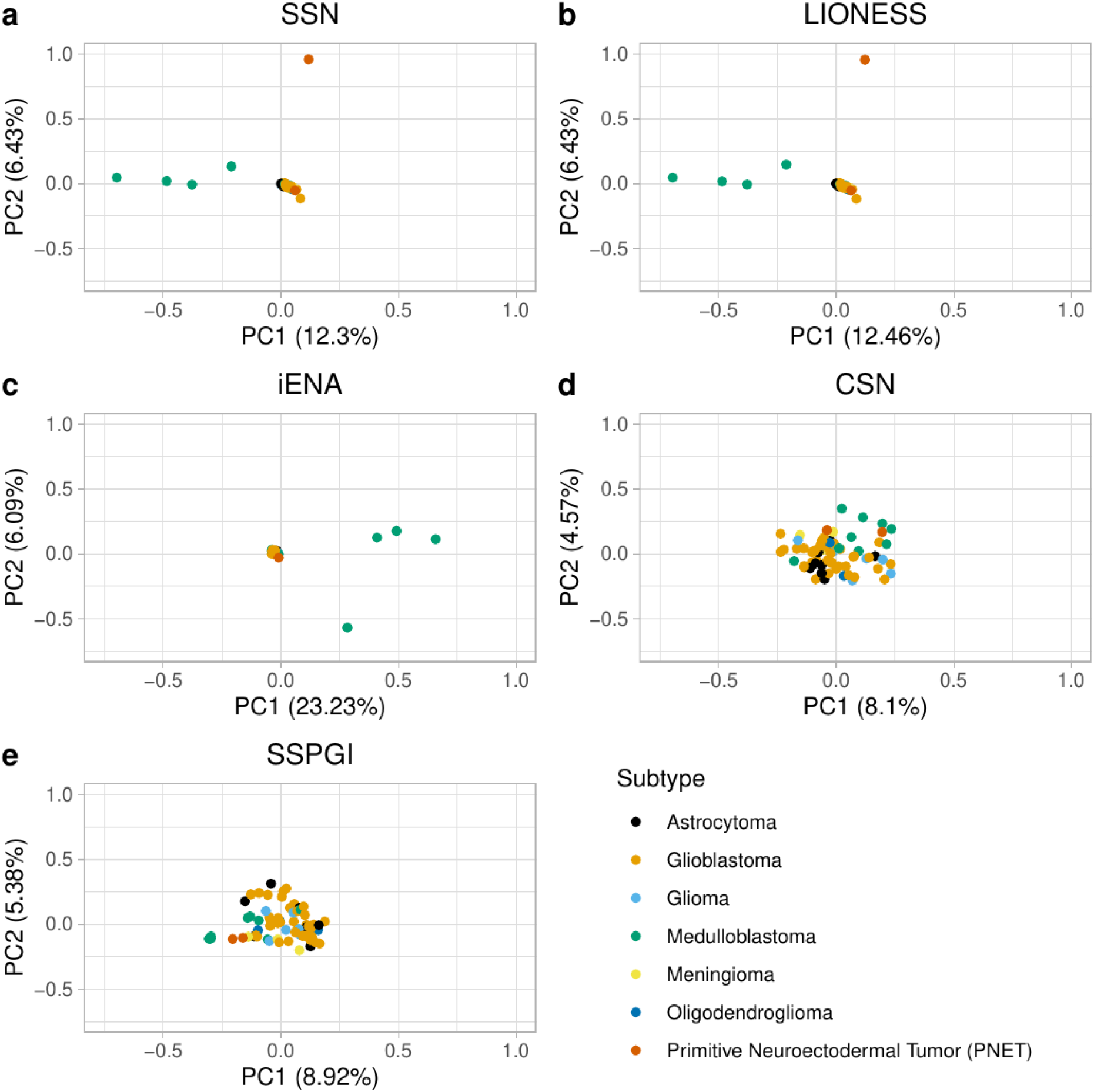
Visualization of brain samples after projecting edge weights onto their first two principal components. We performed PCA analysis on top 25k brain networks constructed using SSN (**a**), LIONESS (**b**), iENA (**c**), CSN (**d**) and SSPGI (**e**). Each dot represents one single-sample network constructed from a cell line corresponding to a given cancer subtype. (PCA: principal component analysis)

**Suppl. Figure 3.**
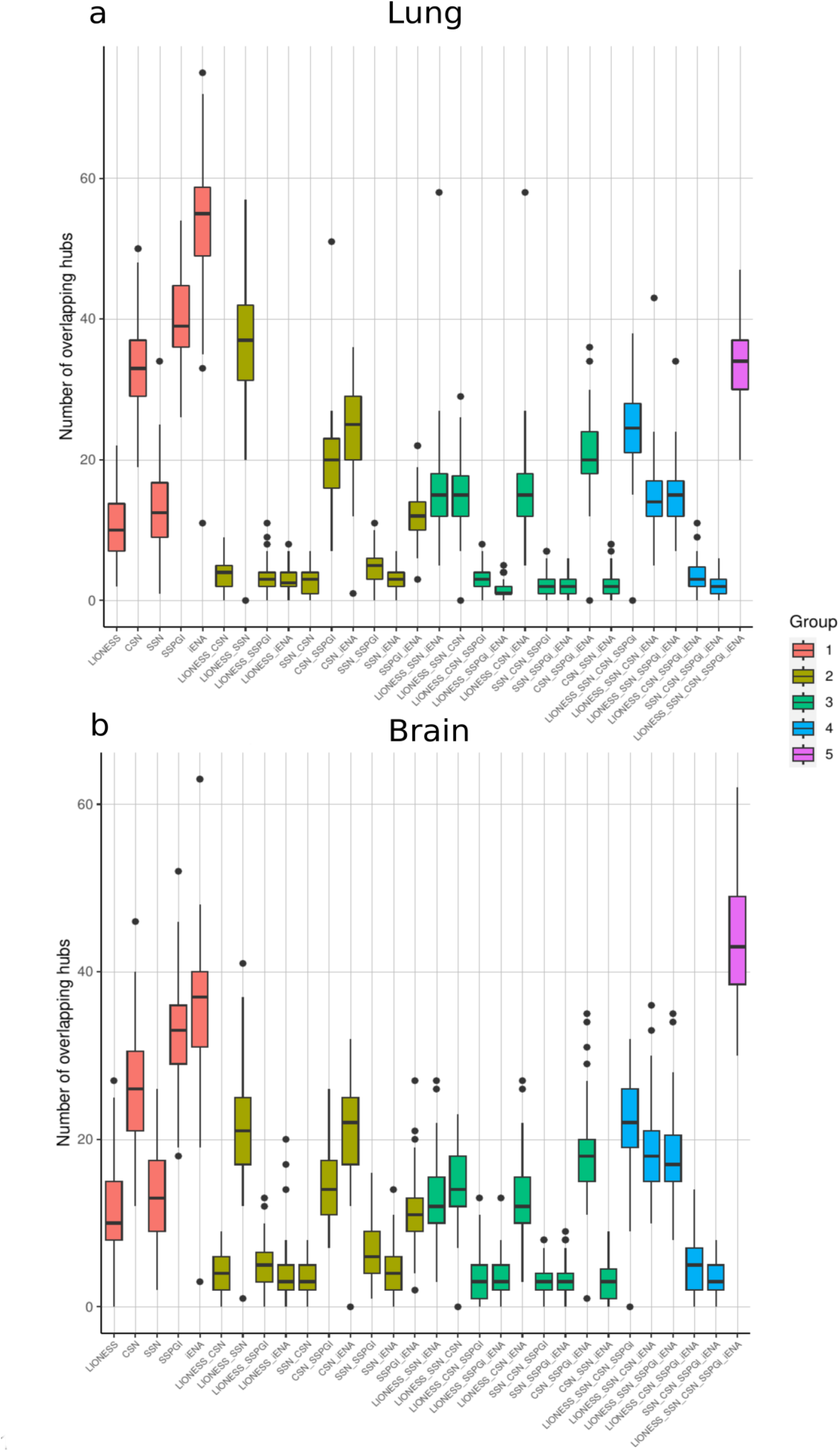
Different genes are identified as hubs in single-sample networks constructed by different methods. For the top 200 nodes of every lung (A) and brain (B) single-sample network, we determined the within-sample overlap across network inference methods. Boxplots represent the average number of overlapping hub genes between 1 (orange), 2 (kaki-green), 3 (green), 4 (blue) or 5 (magenta) distinct single-sample network inference methods.

**Suppl. Figure 4.**
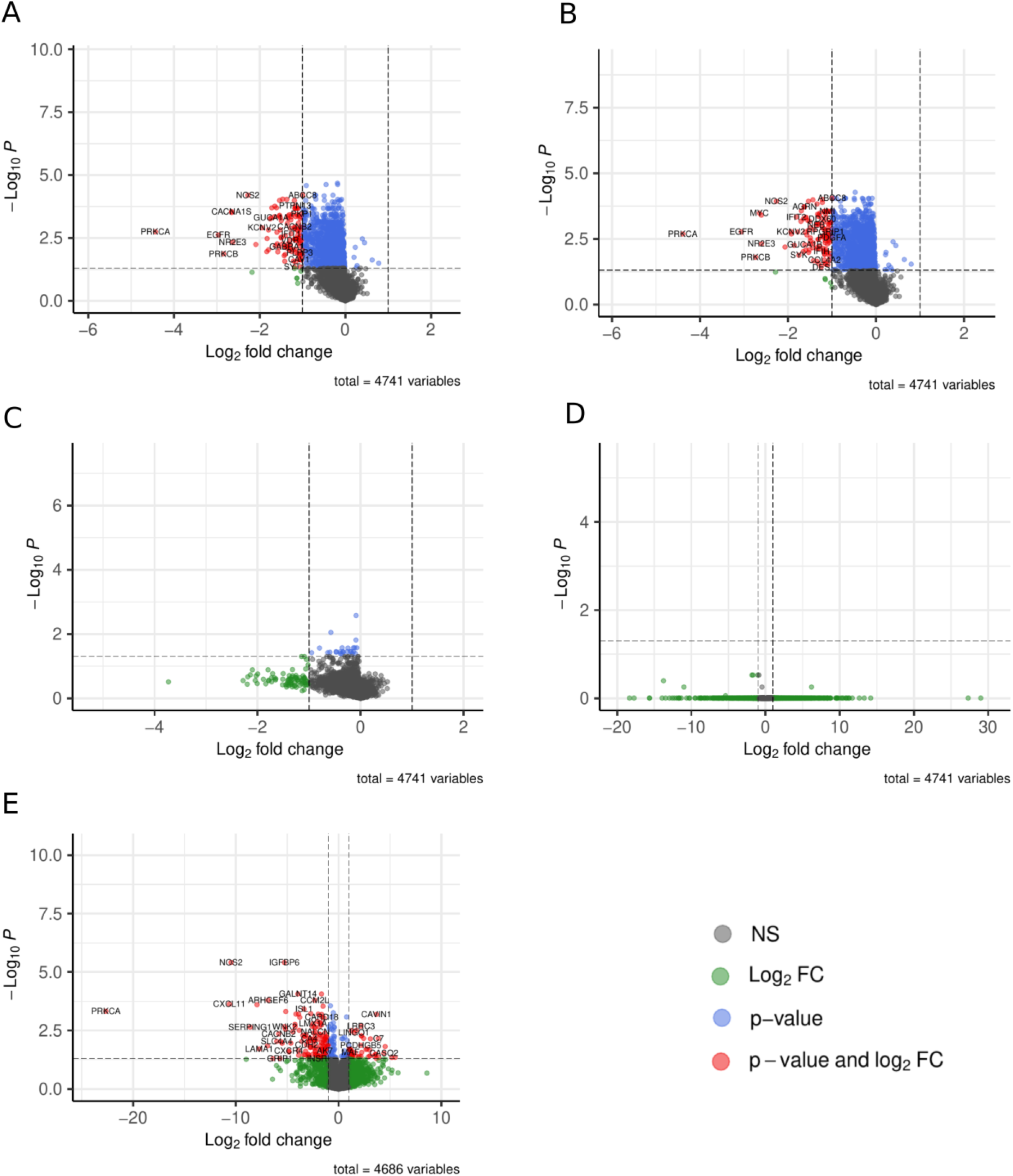
Single-sample networks display distinct differential node importance across network types. The node importance, or sum of absolute edge weights, was calculated for all nodes in top 25k networks constructed by SSN (A), LIONESS (B), iENA (C), CSN (D) and SSPG (E)I. Differentially important nodes (p-adj < 0.05 & |LFC| >= 1) in glioblastoma versus medulloblastoma were identified using linear modelling and an empirical Bayes procedure. (p-adj: adjusted p-value; LFC: log fold change, NS: non-significant)

**Suppl. Figure 5.**
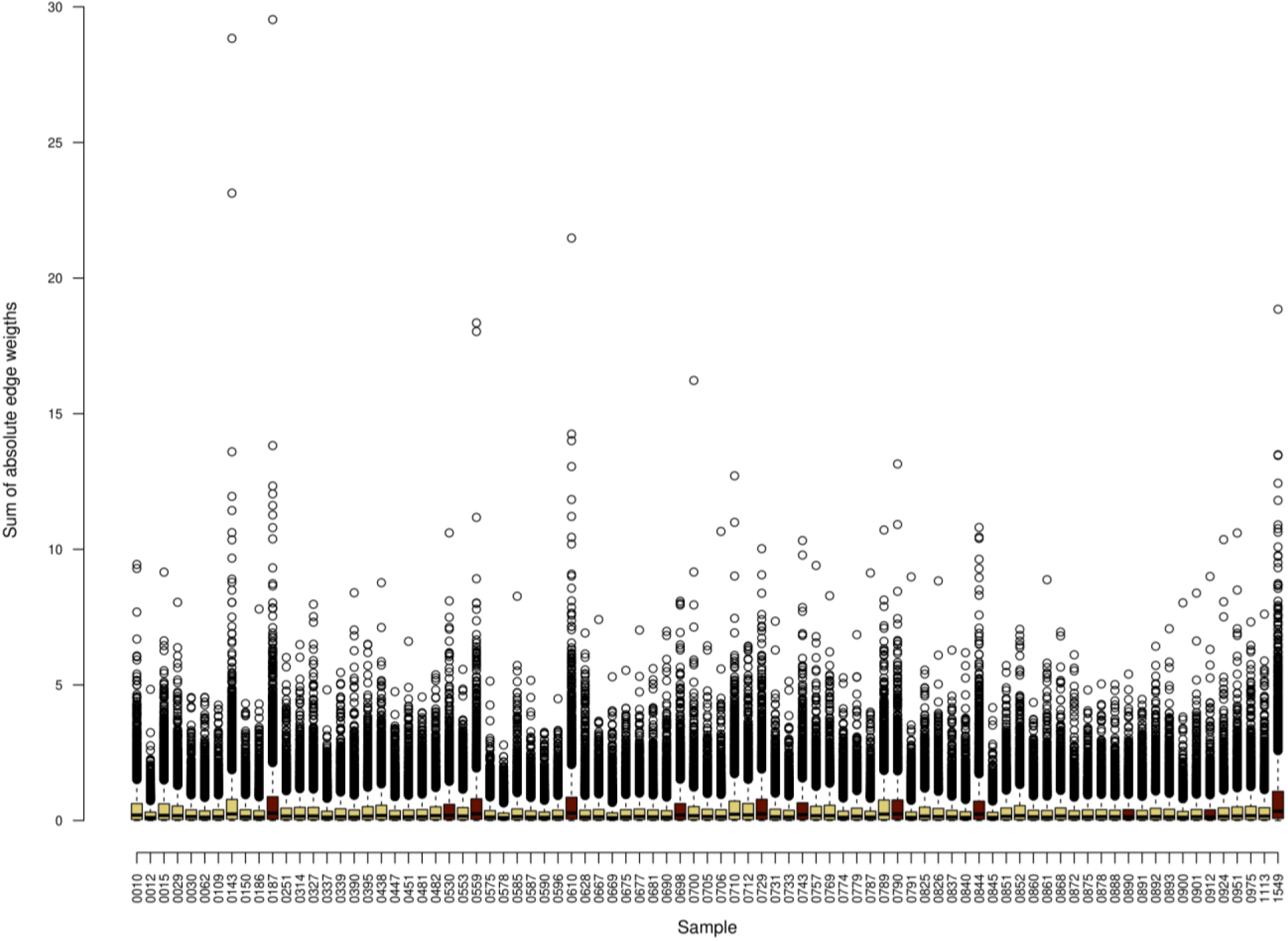
LIONESS lung single-sample networks displayed distinct sum of absolute weight distributions between small cell lung carcinomas (n = 12, wine-red) and non-small cell lung carcinoma (n = 73, green-yellow); The sum of absolute edge weights was calculated for all nodes in top 25k networks constructed by LIONESS.

**Suppl. Figure 6.**
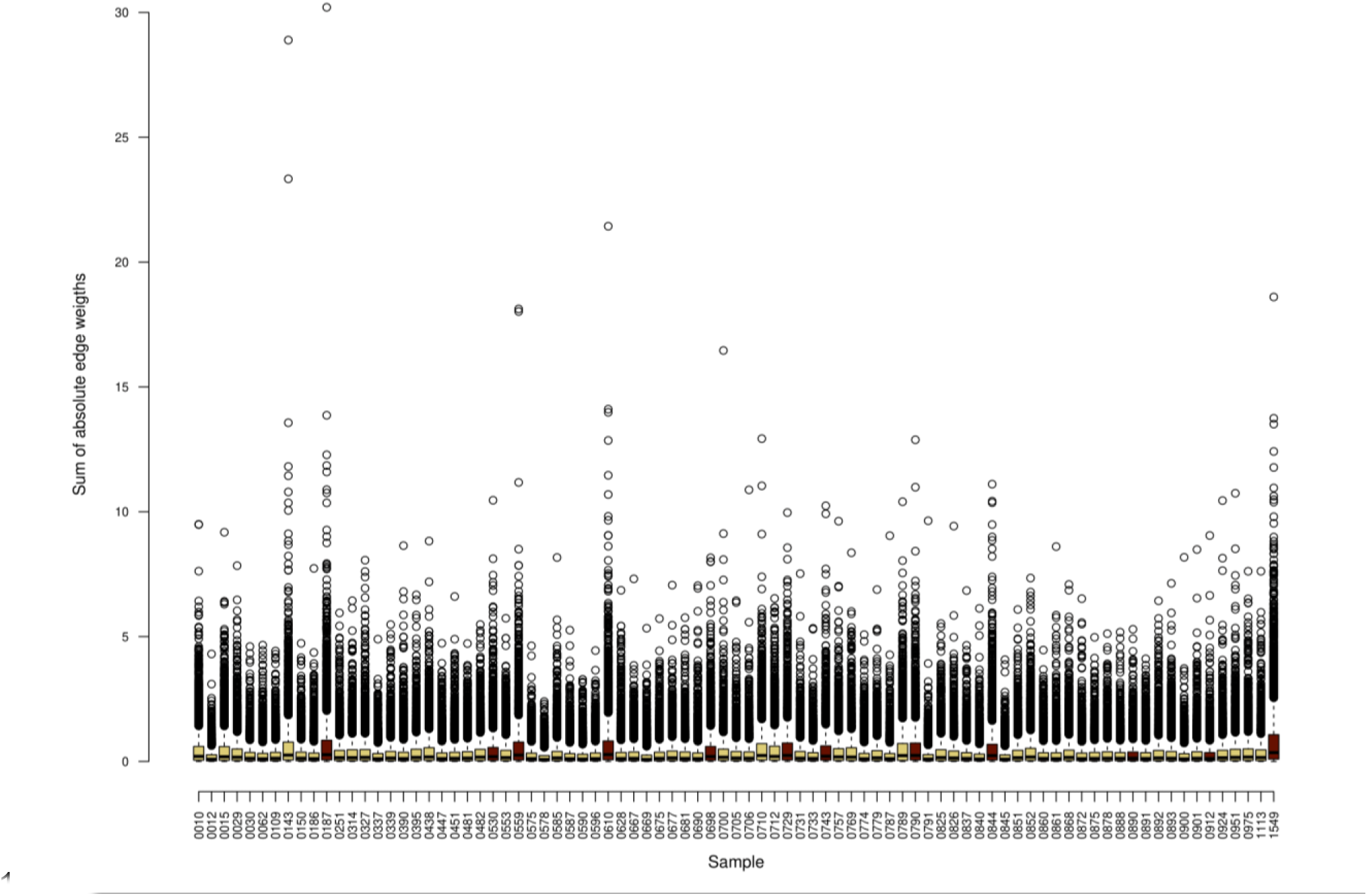
SSN lung single-sample networks displayed distinct sum of absolute weight distributions between small cell lung carcinomas (n = 12, wine-red) and non-small cell lung carcinoma (n = 73, green-yellow); The sum of absolute edge weights was calculated for all nodes in top 25k networks constructed by SSN.

**Suppl. Figure 7.**
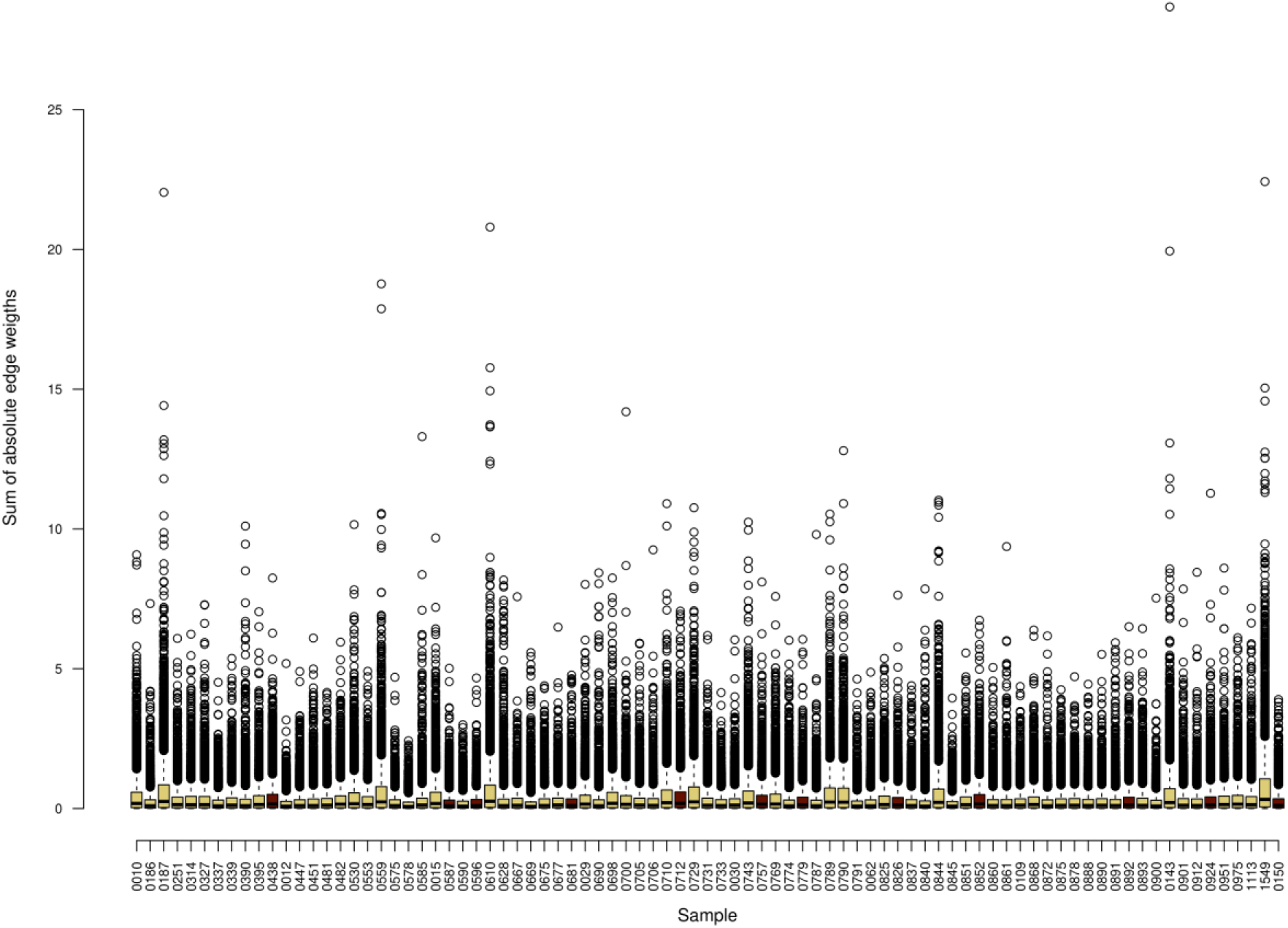
iENA lung single-sample networks did not display distinct sum of absolute weight distributions between small cell lung carcinomas (n = 12, wine-red) and non-small cell lung carcinoma(n = 73, green-yellow); The sum of absolute edge weights was calculated for all nodes in top 25k networks constructed by iENA.

**Suppl. Figure 8.**
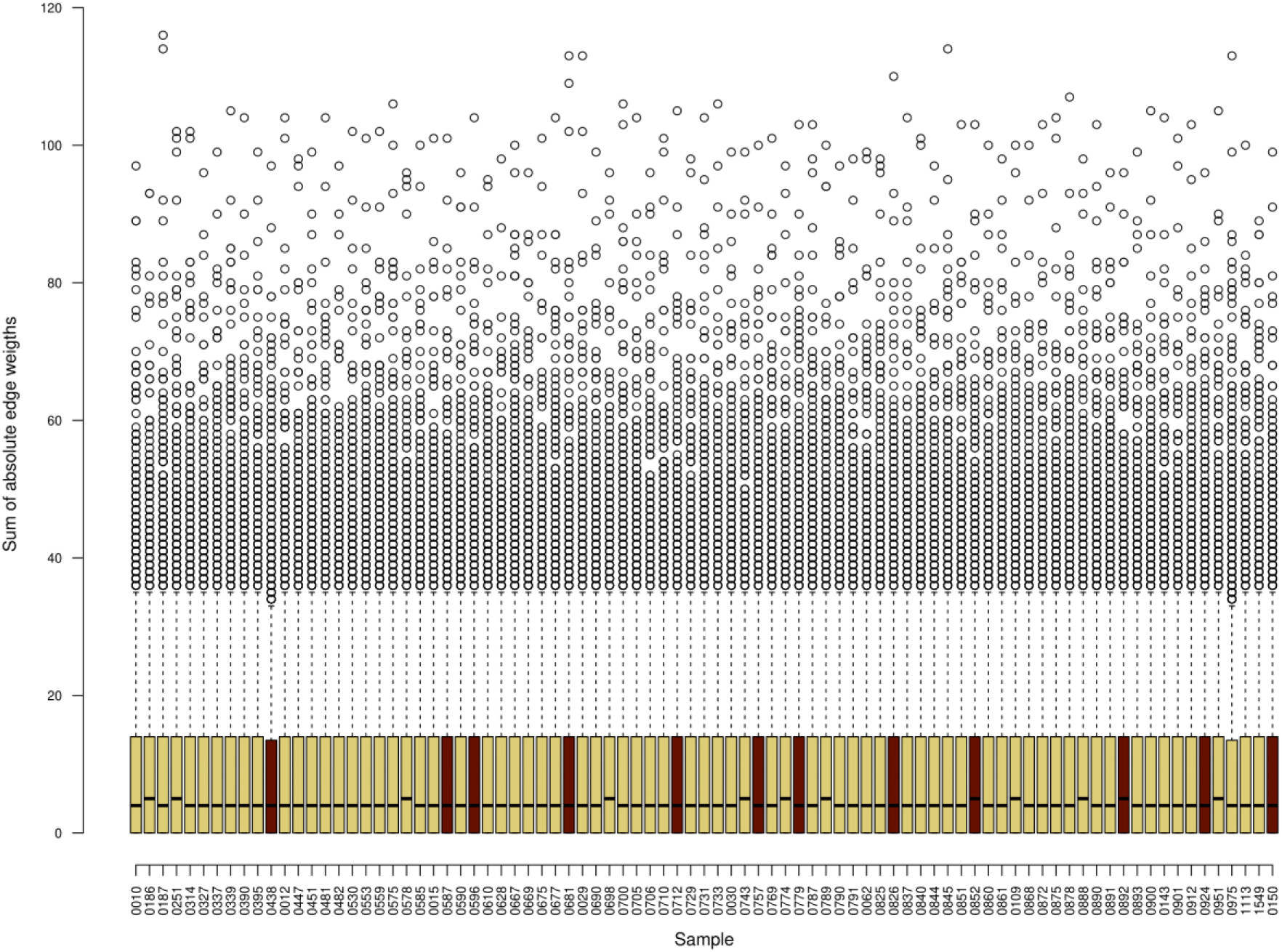
CSN lung single-sample networks did not display distinct sum of absolute weight distributions between small cell lung carcinomas (n = 12, wine-red) and non-small cell lung carcinoma(n = 73, green-yellow); The sum of absolute edge weights was calculated for all nodes in top 25k networks constructed by CSN.

**Suppl. Figure 9.**
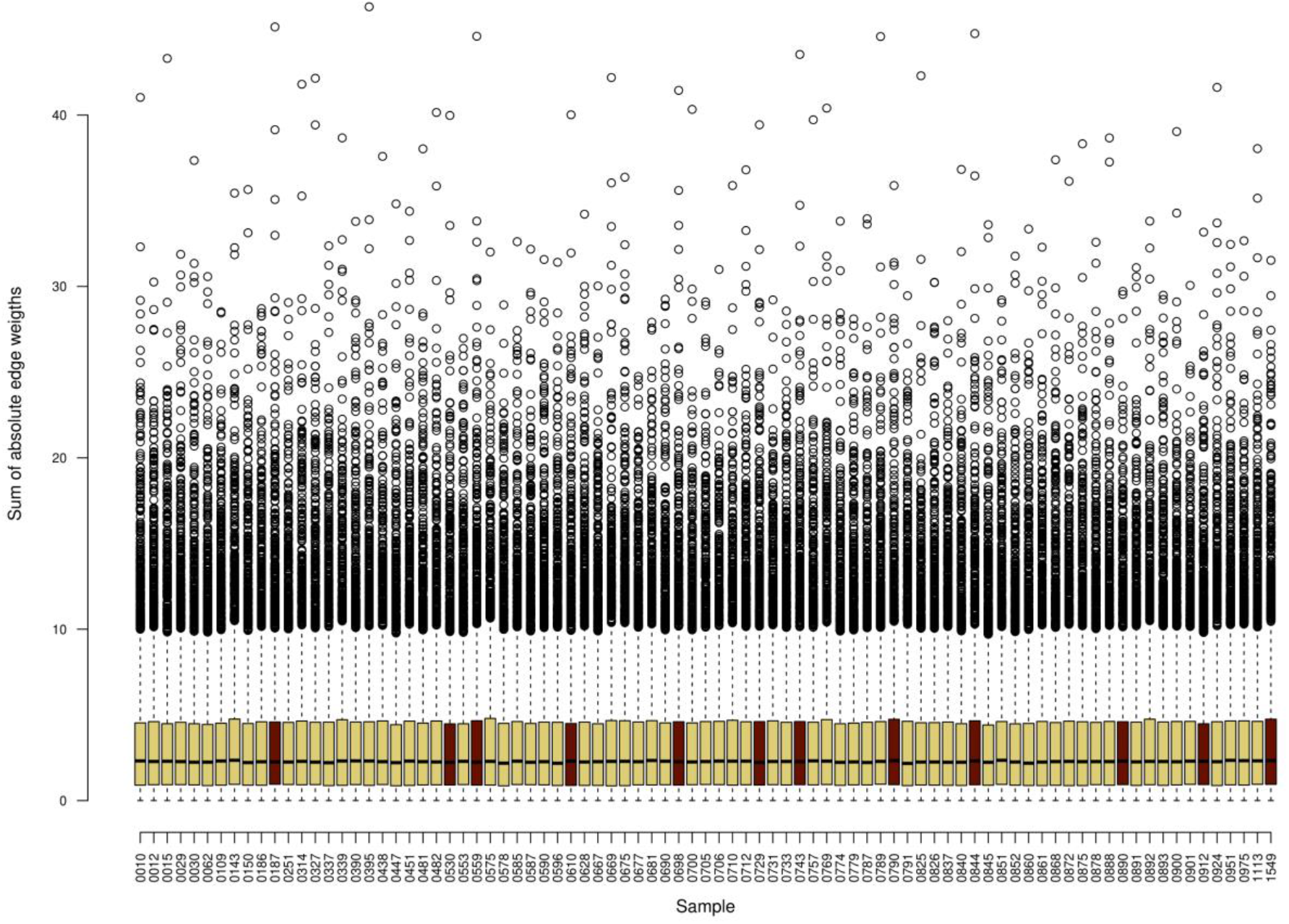
SSPGI lung single-sample networks did not display distinct sum of absolute weight distributions between small cell lung carcinomas (n = 12, wine-red) and non-small cell lung carcinoma(n = 73, green-yellow); The sum of absolute edge weights was calculated for all nodes in top 25k networks constructed by SSPGI.

**Suppl. Figure 10.**
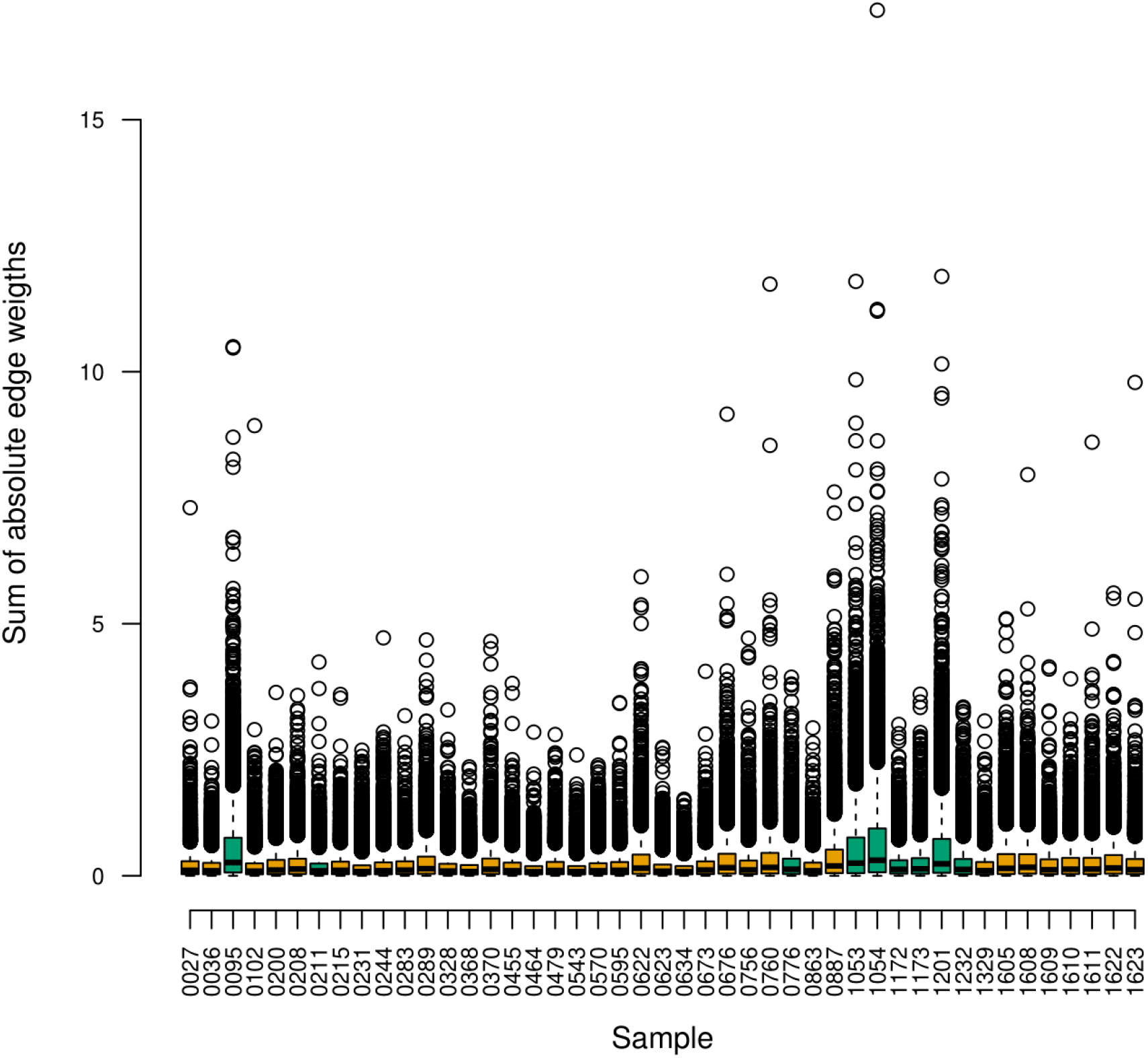
SSPGI brain single-sample networks displayed distinct sum of absolute weight distributions between glioblastoma (n = 53, yellow-orange) and medulloblastoma samples (n = 9, green); The sum of absolute edge weights was calculated for all nodes in top 25k networks constructed by SSPGI.

**Suppl. Figure 11.**
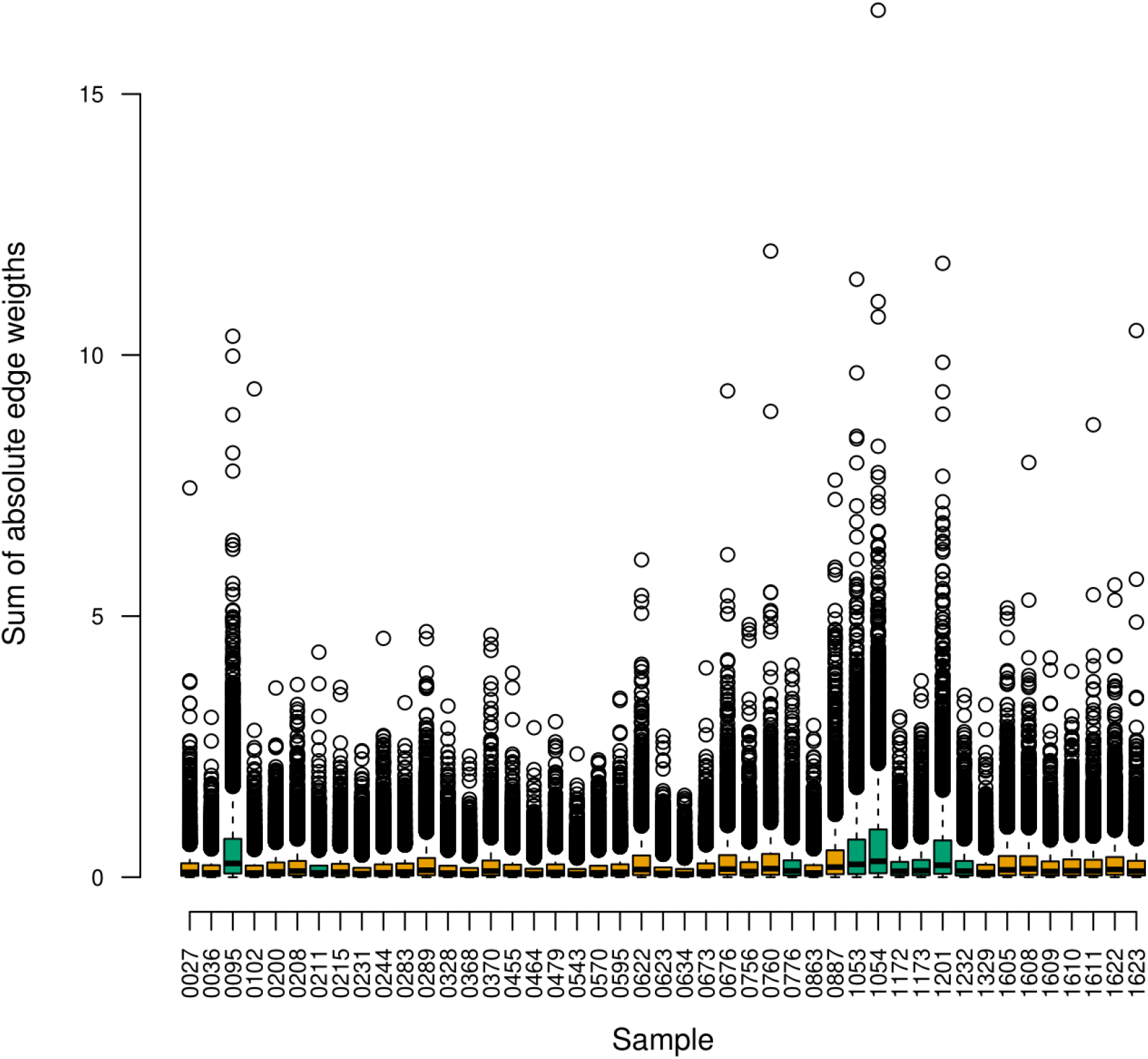
SSN brain single-sample networks displayed distinct sum of absolute weight distributions between glioblastoma (n = 53, yellow-orange) and medulloblastoma samples (n = 9, green); The sum of absolute edge weights was calculated for all nodes in top 25k networks constructed by SSN.

**Suppl. Figure 12.**
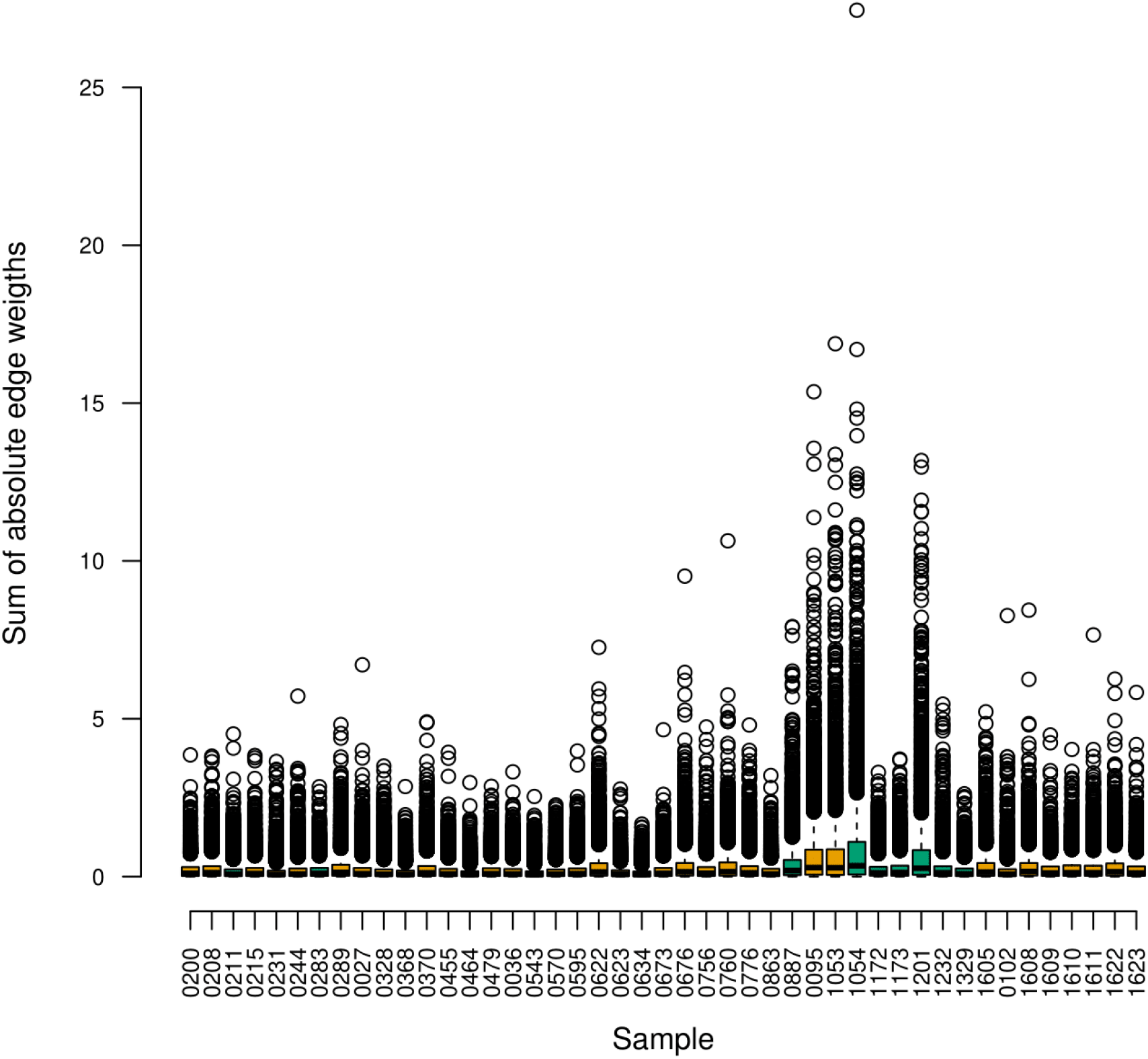
iENA brain single-sample networks displayed distinct sum of absolute weight distributions between glioblastoma (n = 53, yellow-orange) and medulloblastoma samples (n = 9, green); The sum of absolute edge weights was calculated for all nodes in top 25k networks constructed by iENA.

**Suppl. Figure 13.**
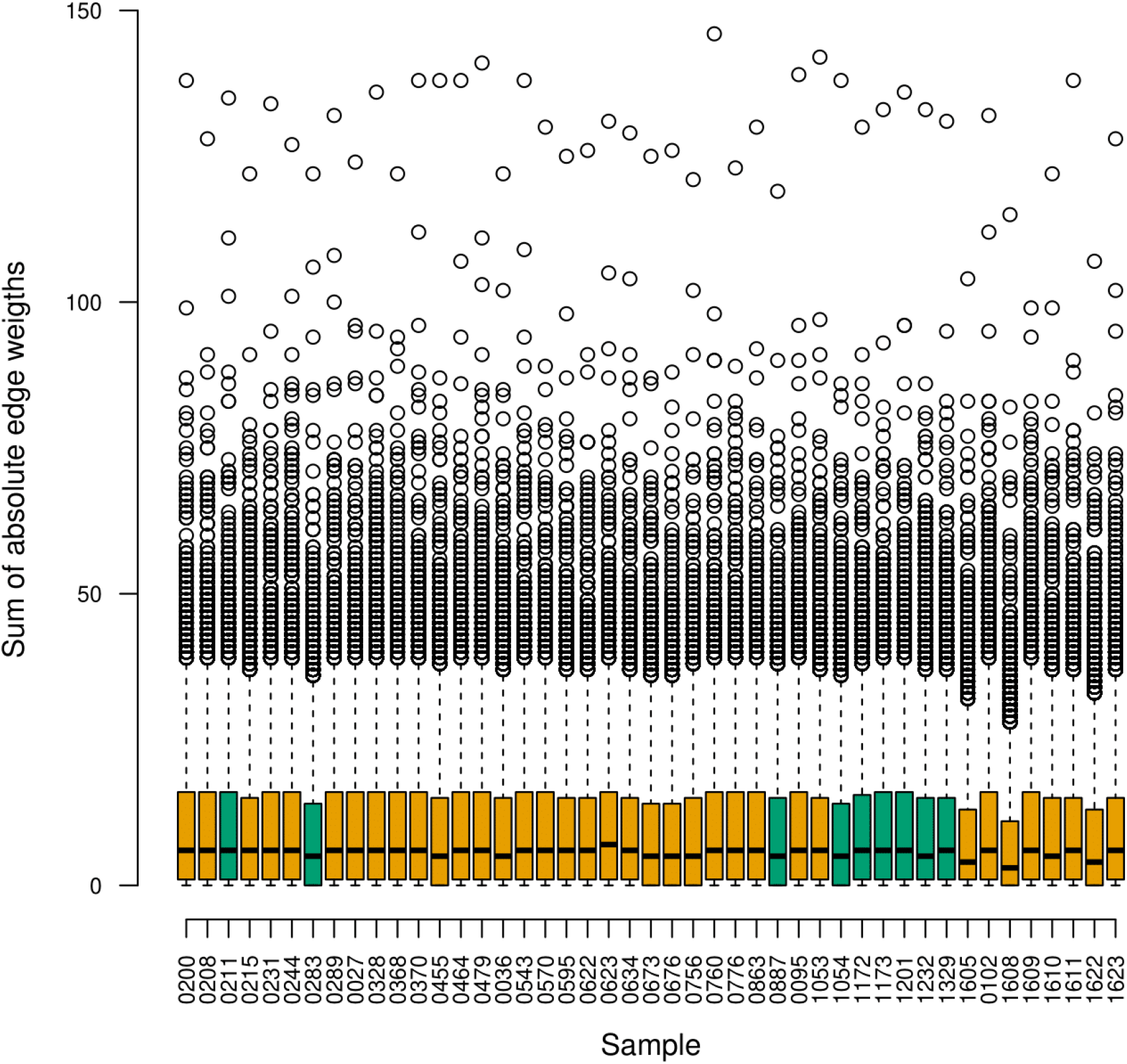
CSN brain single-sample networks did not display distinct sum of absolute weight distributions between glioblastoma (n = 53, yellow-orange) and medulloblastoma samples (n = 9, green); The sum of absolute edge weights was calculated for all nodes in top 25k networks constructed by CSN.

**Suppl. Figure 14.**
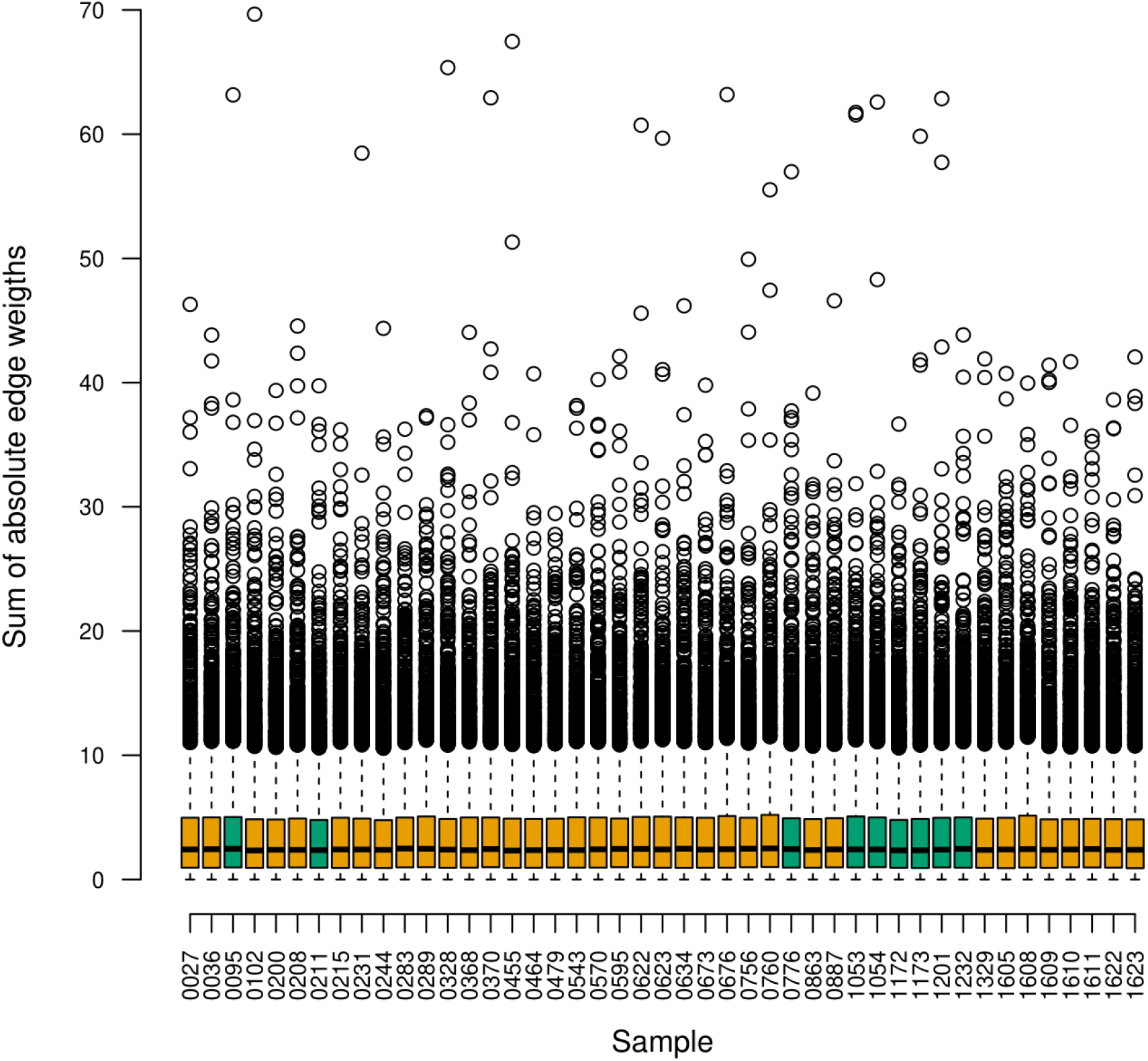
SSPGI brain single-sample networks did not display distinct sum of absolute weight distributions between glioblastoma (n = 53, yellow-orange) and medulloblastoma samples (n = 9, green); The sum of absolute edge weights was calculated for all nodes in top 25k networks constructed by SSPGI.

